# Peripheral Neuronal Activation Shapes the Microbiome and Alters Gut Physiology

**DOI:** 10.1101/2021.04.12.439539

**Authors:** Jessica A. Griffiths, Bryan B. Yoo, Peter Thuy-Boun, Victor Cantu, Kelly Weldon, Collin Challis, Michael J. Sweredoski, Ken Y. Chan, Taren M. Thron, Gil Sharon, Annie Moradian, Gregory Humphrey, Qiyun Zhu, Justin Shaffer, Dennis W. Wolan, Pieter C. Dorrestein, Rob Knight, Viviana Gradinaru, Sarkis K. Mazmanian

## Abstract

The gastrointestinal (GI) tract is extensively innervated by intrinsic neurons of the enteric nervous system (ENS) and extrinsic neurons of the central nervous system and peripheral ganglia, which together regulate gut physiology. The GI tract also harbors a diverse microbiome, but interactions between the ENS and the microbiome remain poorly understood. Herein, we activate choline acetyltransferase (ChAT)-expressing or tyrosine hydroxylase (TH)-expressing gut-associated neurons in mice to determine effects on intestinal microbial communities and their metabolites, as well as on host physiology. The resulting multi-omics datasets support broad roles for discrete peripheral neuronal subtypes in shaping microbiome structure, including modulating bile acid profiles and fungal colonization. Physiologically, activation of either ChAT^+^ or TH^+^ neurons increases fecal output, while only ChAT^+^ activation results in increased colonic migrating motor complexes and diarrhea-like fluid secretion. These findings suggest that specific subsets of peripherally-activated ENS neurons differentially regulate the gut microbiome and GI physiology in mice, without involvement of signals from the brain.

## INTRODUCTION

Diverse cell types in the gastrointestinal (GI) tract coordinate physiology within the gut^1^ and throughout the body^2^. The mammalian gut receives and transmits neuronal signals through ∼100,000 extrinsic nerve fibers originating from the sympathetic, parasympathetic, and sensory nervous systems^3^. The GI tract is also innervated by an extensive network of over 100 million intrinsic neurons organized into two distinct compartments within the GI tract, namely the myenteric plexus and submucosal plexus^4^. The neurons of the GI tract, composing the enteric nervous system (ENS), have been implicated in processes including digestion^5^, immunity^6,7^, and even complex behaviors^8^, in mice. Interactions between neurons of the GI tract and other cell types highlight the diverse roles of the ENS. For example, neuronal pathways in the gut regulate nutrient sensation through intestinal enteroendocrine cells^9^, modulate the epithelial barrier and mucosal immunity^10–12^, and dynamically interface with the microbiome^13,14^. Exposure of the ENS to changing diet, microbiome, and xenobiotics creates inputs distinct from those in the central nervous system (CNS), i.e., the brain and spinal cord.

Choline acetyltransferase (ChAT) and tyrosine hydroxylase (TH) are the rate-limiting enzymes in acetylcholine and catecholamine biosynthesis, respectively, and are key chemical mediators of neurotransmission in the brain and the periphery. Acetylcholine is the primary excitatory neurotransmitter of the gut, and cholinergic neurons represent 60% of the ENS, mediating intestinal propulsion and secretion^15,16^. Several studies have established correlations between neuronal activity, abundance, and specific physiological outcomes^17–19^. For example, age-associated reduction of ChAT^+^ neurons in the ENS coincides with constipation and evacuation disorders^20,21^, and clinical studies have shown that anticholinergic drugs cause constipation and cholinergic agonists can cause diarrhea^22,23^. In a disease context, cholera toxin induces hypersecretion and sustained activation of submucosal ChAT^+^ neurons in mice^24,25^. Although less characterized, TH^+^ neurons and dopamine signaling pathways have also been shown to affect GI motility^26^, and TH^+^ neuronal damage in individuals with Parkinson’s disease (PD) correlates with increased constipation^27,28^.

Though known to be important for motility and secretomotor function, ChAT^+^ and TH^+^ neurons have not yet been systematically characterized and interrogated for their roles in GI physiology^17,29^. One barrier to modulation of neuronal populations in the ENS is its size: 35-40 cm in mice. To circumvent the need for direct delivery of effectors, we leveraged a systemically-delivered engineered adeno-associated virus (AAV) with enhanced tropism for the ENS and other peripheral ganglia of mice ^30^. Importantly, this vector, AAV-PHP.S, does not transduce the CNS, allowing us to uncouple peripheral activation from brain-to-gut signaling. We find that activating gut-associated ChAT^+^ and TH^+^ neurons of mice with chemogenetic modulators^31^ alters the transcriptional and proteomic landscape of the intestines, as well as the gut metagenome and metabolome. Multi-‘omic’ analyses allow us to characterize detailed and complex host-microbial interactions, and enable prediction of neuronal influences on a number of biological processes in the gut, including providing insights into secondary bile acid production and control of fungal populations, among other interesting associations. In addition, we show that activation of gut-associated neurons strikingly impacts GI function, including motility and fluid secretion. Together, this work reveals differential effects of non-brain activation of ChAT^+^ and TH^+^ neurons in shaping the gut environment and GI physiology and generates rich datasets as a resource for further exploration (DOI:10.5281/zenodo.10525220, https://github.com/mazmanianlab/Griffiths_Yoo_et_al/).

## RESULTS

### Distinct spatial localization of ChAT^+^ and TH^+^ neurons in the ENS

Broad ENS morphology has been previously characterized using immunohistochemistry (IHC)^21,32,33^. To map neurons in mice with higher resolution, we used recombinant AAVs to fluorescently label enteric neurons *in vivo*, and tissue clearing techniques to enhance visualization of intact GI tissue^34–36^. Imaging whole tissue, without the need for sectioning, preserves neuronal architectures over large distances and across both longitudinal and cross-sectional axes. The AAV capsid variant AAV-PHP.S is optimized for systemic delivery in mice^37^ and displays increased tropism for the peripheral nervous system (PNS), including the ENS^38^. To further optimize ENS expression, we replaced the CAG promoter used in^30^ with the human Synapsin 1 (hSYN1) promoter, which has been shown to restrict gene expression to neurons^38^ and minimize expression in peripheral targets such as the dorsal root ganglia (DRGs)^39^. To assess off-target effects, we compared expression of AAV-PHP.S-delivered hSYN1-mNeonGreen to that of CAG-mNeonGreen in various non-ENS tissues known to affect GI function (Figure S1). Expression from the hSYN1 construct only sparsely labeled the DRGs and jugular-nodose ganglia, and did not label neuronal projections in the vagus nerve or dorsal root, unlike the previously-used CAG construct (Figures S2A and S2B)^30^. In the CNS, AAV-PHP.S-hSYN1 did not label neurons in the brain, brainstem, or spinal cord (Figure S3).

We packaged genes encoding fluorescent proteins (tdTomato or mNeonGreen) under control of the hSYN1 promoter into AAV-PHP.S, delivered them systemically, and found that 90% (±2.6% SD) of ENS cells labelled with antibodies against *<u>P</u>*rotein *<u>G</u>*ene *<u>P</u>*roduct 9.5 (PGP9.5), a pan-neuronal protein, co-localized with virally-labelled neurons in the small intestine (SI) and colon (Figure 1A). A single systemic injection of AAV-PHP.S-hSYN1-mNeonGreen at a dose of 10^12^ viral genomes (vg) was sufficient to label spatially diverse regions of the ENS, such as ganglia proximal and distal to the mesentery (Figure S4A). Viral transduction was uniform throughout the SI and colon, aside from a small (∼1.5 cm) section of the medial colon that, for unknown reasons, was consistently not well transduced and was therefore excluded from further analysis (Figure S4B).

**Figure 1.**
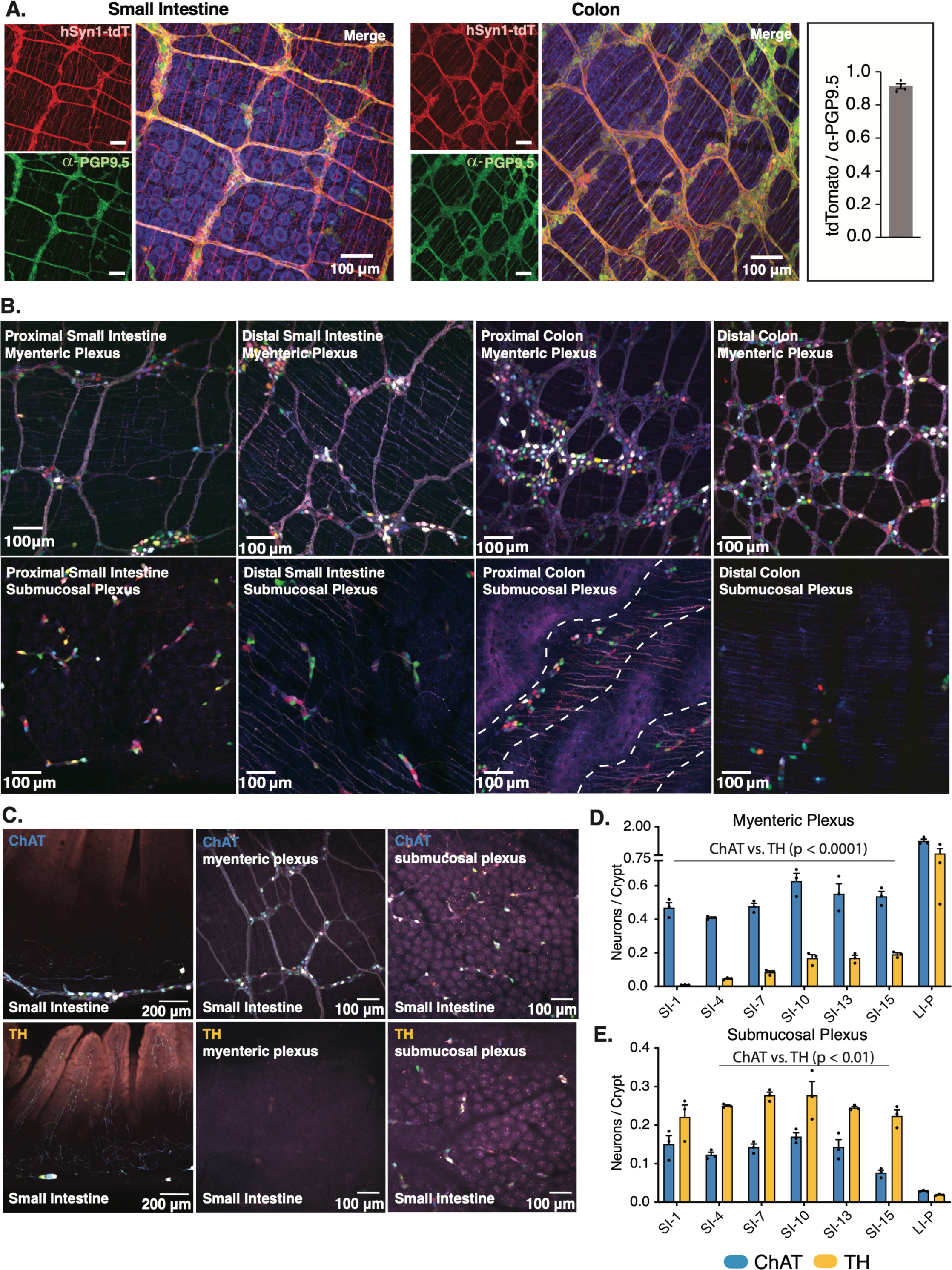
ChAT^+^ versus TH^+^ Neuronal Distribution in the ENS. (A) Representative images of SI and colon from mice infected with AAV-PHP.S-hSYN1-tdTomato and immunolabelled with the pan-neuronal antibody PGP9.5. Inset shows quantification of the ratio of tdTomato^+^ cells / PGP9.5^+^ cells (N=3 mice, each data point represents the average of 3 representative fields). (B) Representative images of proximal and distal regions of the SI and colon from AAV-PHP.S-hSYN1-XFP infected mice. Dotted lines demarcate the rugae (folds) in the proximal colon. (C) Representative images of cross-sections, myenteric, and submucosal plexuses in ChAT-Cre and TH-Cre mice infected with AAV-PHP.S-hSYN1-DIO-XFP. (D-E) Density of neurons in the myenteric plexus and submucosal plexus of ChAT-Cre and TH-Cre mice normalized to the number of crypts (N=3 mice, each data point represents the average of 3 representative fields ChAT-Cre vs. TH-Cre mice were compared using two-way ANOVAs with Sidak’s correction for multiple comparisons for the SI and LI independently; Comparison of different regions in the SI of ChAT-Cre or TH-Cre mice were analyzed using one-way ANOVAs with Tukey’s correction for multiple comparisons). See also Figures S1-S5 and Video Supplement 1 **Source Data** Figure 1 https://github.com/mazmanianlab/Griffiths_Yoo_et_al/tree/main/ENS%20quantification

To explore the general architecture of the ENS, we transduced wild-type mice with a single i.v. injection of a pool of AAV-PHP.S packaging multiple fluorescent proteins (AAV-PHP.S-hSYN1-XFP), which broadly labelled enteric neurons in the gut and enabled us to distinguish cells by distinct colors resulting from stochastic transduction with different combinations of XFPs (Figure 1B). We quantified the number of neurons and ganglia, as well as the ganglion size (i.e., the number of neurons in each ganglion) in the myenteric and submucosal plexuses of seven regions of the SI and two regions of the colon (Figures S5A-S5F). Regions were approximately 1 cm in length and the tissue was sampled every 2-3 cm. We saw that in the SI, the numbers of neurons and ganglia generally increased toward the distal portion of the myenteric plexus, while the converse was true for the submucosal plexus (i.e., lower numbers in distal than proximal regions) (Figures S5A and S5B). Additionally, the size of the ganglia (i.e., the number of neurons per ganglion) increased in the distal region of the SI myenteric plexus, a feature not observed in the submucosal plexus (Figure S5C). While neuronal numbers were similar in the proximal and distal regions of the colonic plexuses (Figure S5D), the number of myenteric ganglia increased (Figure S5E) while the size of each ganglion decreased in the distal colon (Figure S5F). Interestingly, submucosal neurons in the proximal colon localized to natural folds in the tissue (Figure 1B, dashed lines in lower second-from-right panel).

To visualize ChAT^+^ and TH^+^ neurons, we employed mouse lines in which Cre recombinase (Cre) is expressed under the control of the respective gene promoter and engineered viral constructs with the transgene in a double-floxed inverted orientation (DIO) so that the transgene is flipped and expressed in a Cre-dependent manner. After transducing ChAT-Cre or TH-Cre mice with AAV-PHP.S-hSYN1-DIO-XFP, we observed that both neuronal populations occupy spatially distinct layers of the GI tract, with ChAT^+^ neurons primarily located in the myenteric plexus and TH^+^ neurons more abundant in the submucosal plexus (Figure 1C). Quantifying this effect, we found more ChAT^+^ than TH^+^ neurons in all assayed regions of the myenteric plexus (Figure 1D), although the density of TH^+^ myenteric neurons increased distally (Figure 1D, 10-fold increase from SI-1 vs SI-7; 2-fold increase from SI-7 vs SI-10/13/15). In the small intestine, by contrast, there were more TH^+^ than ChAT^+^ submucosal neurons (Figure 1E). In addition to providing these insights into ENS architecture, this approach for whole tissue imaging without the need for antibody labeling (which has limited penetration to deeper layers) should be broadly useful for profiling other neuronal and non-neuronal cell types in the gut.

### Activation of gut-associated neurons reshapes the gut microbiome

The unique spatial organization of ChAT^+^ and TH^+^ neurons we observed suggests potentially distinct functions, which we decided to investigate through specific activation of each neuronal population. First, we examined the specificity of AAV-PHP.S-hSYN1 by staining gut-extrinsic PNS ganglia for TH, and found no transduction of TH^+^ cells in the DRGs or jugular-nodose ganglia (Figures S2B and S2C). ChAT^+^ neurons are absent in these peripheral ganglia (Figure S2C) ^40,41^. Prior research has shown that AAV-PHP.S-hSYN1 transduces the prevertebral sympathetic ganglia, which are known to innervate the gut ^42^, but these ganglia also lack ChAT^+^ neurons ^43^. In fact, the vast majority of neurons in the prevertebral sympathetic ganglia are TH^+^ ^43,44^.Therefore, for the remainder of the manuscript, we will use the term “gut-associated” to refer to ChAT^+^ neurons in the ENS, or TH^+^ neurons in the ENS plus innervating prevertebral sympathetic ganglia.

For cell-specific neuronal activation, we employed a Cre-dependent genetic construct encoding an activating ‘*<u>D</u>*esigner *<u>R</u>*eceptor *<u>E</u>*xclusively *<u>A</u>*ctivated by *<u>D</u>*esigner *<u>D</u>*rugs’ (DREADD), named hM3Dq, which is a modified neurotransmitter receptor designed to induce neuronal activation when exposed to Compound 21 (C21), a “designer drug” specific to this receptor^45^. We validated functional gene delivery and expression using intestinal explants from a ChAT-Cre mouse transduced with the activating DREADD and a construct encoding the calcium sensor GCaMP6f, observing a gradual increase in fluorescence consistent with a calcium transient following administration of C21 (Figure S5G and Video Supplement 1).

We reasoned that neuronal activation in the gut may impact the composition and community structure of the gut microbiome. Accordingly, we transduced ChAT-Cre or TH-Cre mice with either virus carrying the activating DREADD (AAV-PHP.S-hSYN1-DIO-hM3Dq-mRuby2) or a control virus expressing only the fluorescent reporter protein (AAV-PHP.S-hSYN1-DIO-mRuby2). We performed shotgun metagenomics on a longitudinal series of fecal samples collected prior to and following ChAT^+^ or TH^+^ neuron activation by C21 (on days 2, 6, and 10 of C21 administration), as well as contents of the terminal cecum collected on day 10 (Figure 2A). In ChAT^+^-activated mice, Faith’s phylogenetic diversity (i.e., alpha-diversity) decreased dramatically in the day 10 fecal and cecal samples (Figure 2B), with many microbial taxa less abundant (Figures 2I-2K; Figure S6; Figure S7). In contrast, TH^+^-activated mice displayed similar phylogenetic diversity to controls throughout the experiment (Figure 2B). Using weighted UniFrac distances and principal coordinate analysis (PCoA) to determine the composition of microbial communities (i.e., beta-diversity), we observed a distinction between ChAT^+^-activated and control animals in both feces and cecal contents, a shift that was absent in samples from TH^+^-activated mice and controls (Figures 2C-H). Over the experimental time course, Verrucomicrobia became significantly enriched in ChAT^+^-activated mice (Figure 2I). To explore differentially abundant bacterial taxa, we used linear discriminant analysis effect size (LEfSe)^46^ and generated cladograms depicting the phylogenetic relationships of differentially abundant taxa (Figures 2J-2M). This analysis revealed that the bacterial species *Akkermansia muciniphila* drove the increase in Verrucomicrobia we observed in ChAT^+^-activated mice (Figures 2N and 2O).

**Figure 2.**
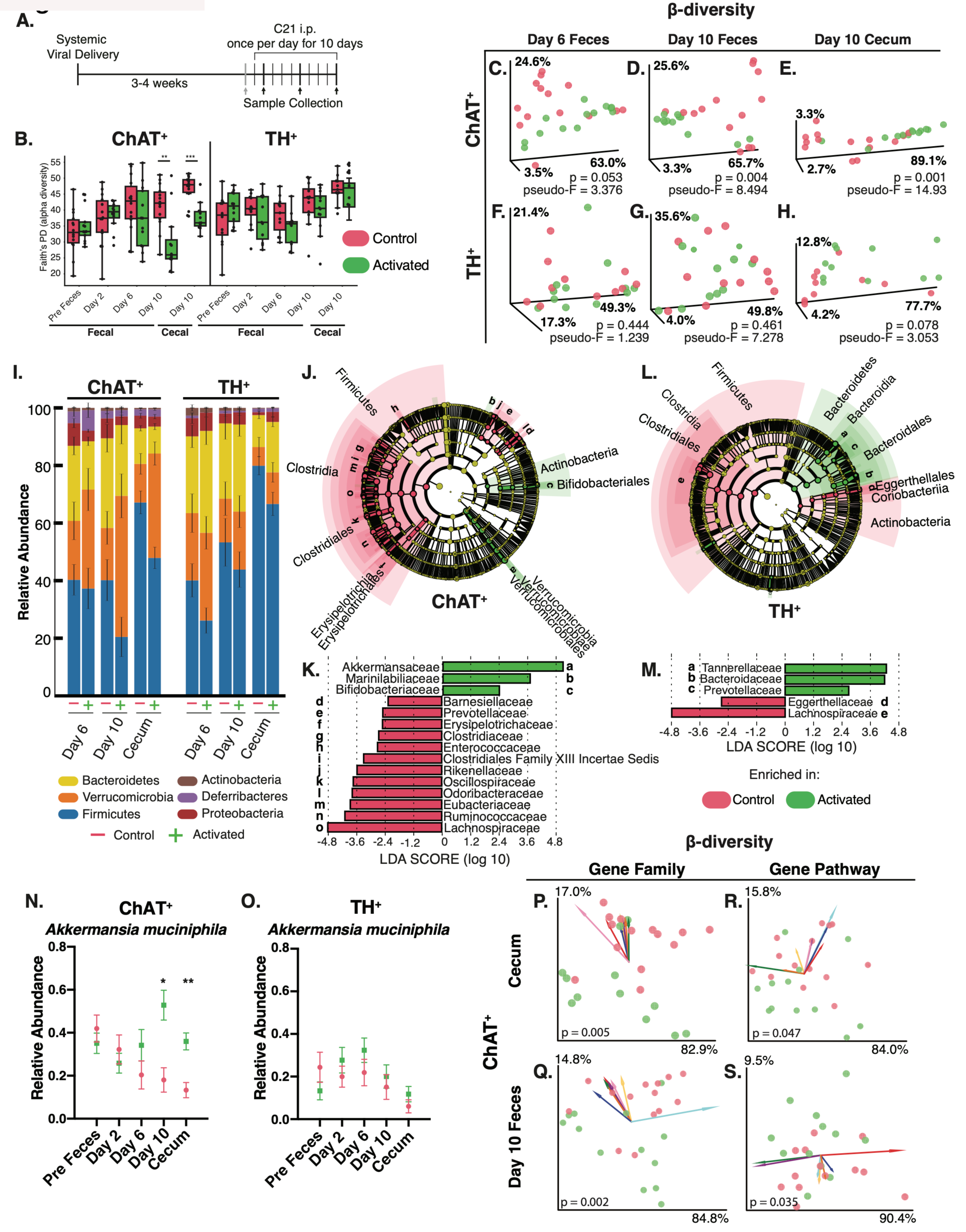
Gut-Associated ChAT^+^ and TH^+^ Neuronal Activation Alters the Gut Microbiome. (A) Experimental paradigm: Cre-dependent hM3Dq was virally administered to either ChAT-Cre or TH-Cre mice. After 3-4 weeks of expression, C21 was injected daily for 10 days to induce specific neuronal activation. Feces were sampled the day prior to the first C21 injection and on days 2, 6, and 10 of C21 administration, and tissue and cecal contents were collected one hour after the last injection. (B) Faith’s phylogenetic diversity of feces and cecal contents over 10 days of neuronal activation in ChAT^+^ and TH^+^ mice. Feces were collected pre-experiment (1 day before first injection) and on day 2, 6, and 10. Cecal contents were collected at experimental endpoint on day 10. (**p<0.01, ***p<0.001 determined by Kruskal-Wallis one-way ANOVA). (C-H) Weighted UniFrac principle coordinate analysis (PCoA) of Activated vs. Control in ChAT^+^ and TH^+^ mice. Statistics performed in QIIME2 as in Bolyen et al., 2019^95^ (I) Stacked bar graph showing phylum-level changes in relative abundance on day 6 and day 10 of injection for feces and day 10 for cecal contents. (J-M) Linear discriminant analysis (LDA) Effect Size (LEfSe) of the cecal microbiome. Cladograms showing differential phylogenetic clusters and family-level differences in activated and control (J,K) ChAT^+^ and (L,M) TH^+^ mice (Cutoff: log_10_(LDA Score) > 2 or < −2) (N-O) Changes in relative abundance of *Akkermansia muciniphila* in feces and cecal contents of (N) ChAT^+^ and (O) TH^+^ mice. (n=11-14 mice per group, per time point; red=Control, green=hM3Dq-Activated, *p<0.05, **p<0.01, determined by multiple t-tests with Holm-Sidak correction for multiple comparisons) (P-S) Beta-diversity of bacterial gene families and pathways in the (P,R) cecum and (Q,S) post-9 feces of control and activated mice. The direction and length of arrows indicate their influence in separating control and activated groups. Colors represent gene families and pathways (annotated in Figure S7). See also Figures S6 and S7 **Source Data** Figure 2 https://github.com/mazmanianlab/Griffiths_Yoo_et_al/tree/main/metagenomics

In addition to identifying microbial species, metagenomic analysis can reveal gene families and pathways that are differentially abundant in the microbiome. ChAT^+^-activated mice, but not TH^+^-activated mice, showed changes in beta-diversity of both gene families and pathways, with shifts evident in the cecal contents and feces collected 9 days after activation (Figures 2P-2S). The most distinguishing features were highly represented in the control group and downregulated in ChAT^+^-activated mice and were mainly associated with bacterial processes, such as nucleotide biosynthesis and metabolism, and protein translation and transport (Figures 2P-2S; Figures S7C and S7D). This downregulation is consistent with the decrease in bacterial alpha-diversity we observed in ChAT^+^-activated mice (Figure 2B). We conclude that neuronal activation actively reshapes the gut microbiome at community, species, and genetic levels, with considerable differences between the effects of ChAT^+^ and TH^+^ neurons.

### Neuronal stimulation impacts the gut metabolome

Given the intimate and intertwined mouse and microbial co-metabolism, the changes in the microbial metagenome we observed in response to neuronal activation led us to predict that there would also be alterations in the profile of gut metabolites. We therefore performed untargeted metabolomics using liquid chromatography with tandem mass spectrometry (LC-MS/MS) to assay molecular changes in cecal contents and feces following neuronal activation in the gut. In both ChAT^+^- and TH^+^-activated neurons, compared to unactivated controls (no DREADD), we observed a strong separation of metabolome profiles in cecal samples taken one hour following the last C21 injection (Figures 3A and 3B). Thus, targeted activation of ChAT^+^ and TH^+^ gut-associated neurons appears to strongly influence the gut metabolome.

**Figure 3.**
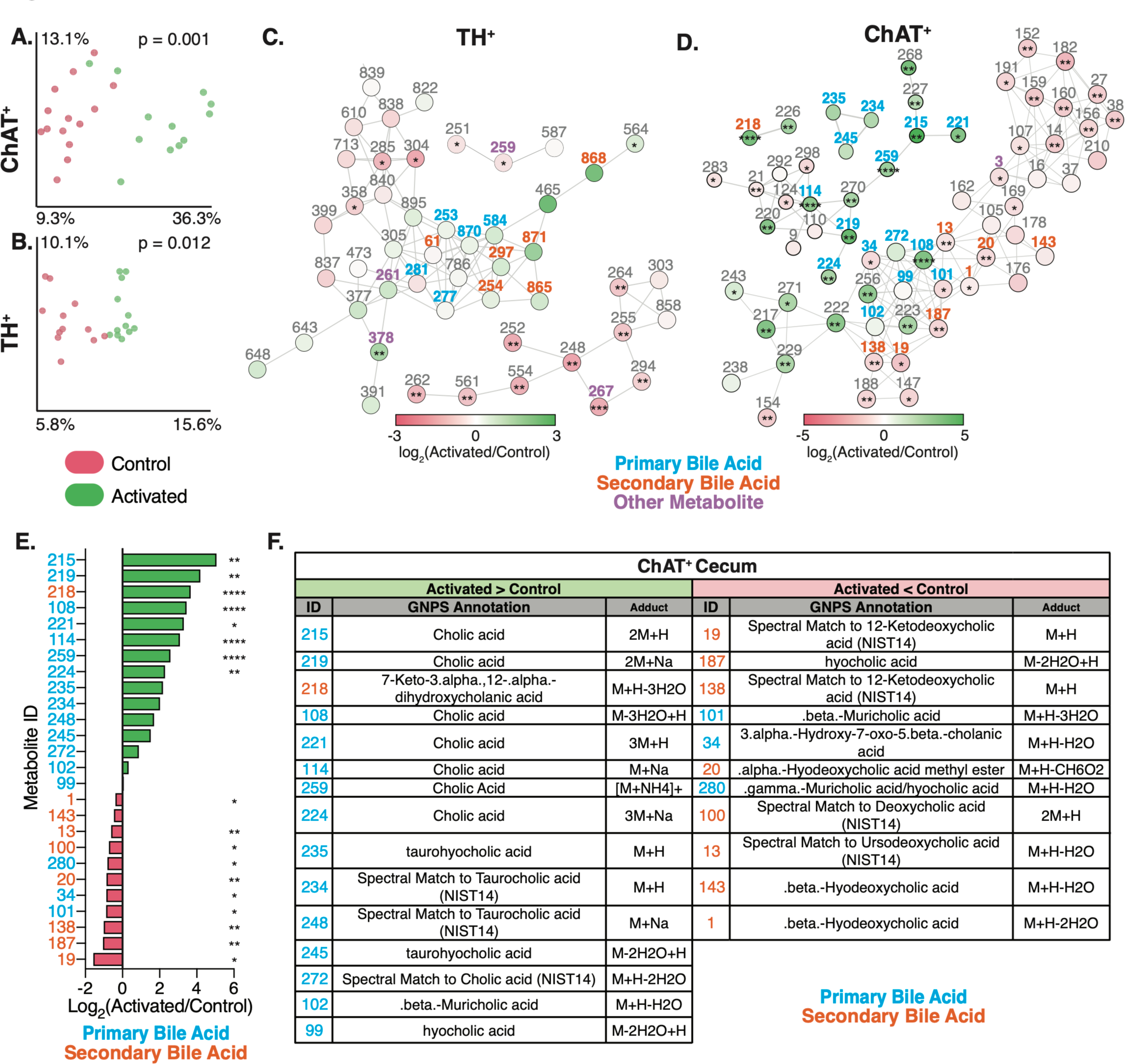
Gut-Associated ChAT^+^ or TH^+^ Neuronal Activation Alters Host and Microbe-Derived Luminal Metabolites. (A-B) Canberra PCoA of the cell-free, luminal metabolome of cecal contents from control (red) and activated (green) ChAT^+^ and TH^+^ mice. Statistical analyses were performed in QIIME2 as in Bolyen et al., 2019^95^ (C-D) Metabolic networks constructed from identified cecal metabolites in TH^+^ and ChAT^+^ mice. Each node is colored by its upregulation (green) or downregulation (red) in the activated group and is labelled with an ID number corresponding to annotation, mass-to-charge ratio, retention time, fold change, and significance value in Table S1 (E) Fold changes of specific bile acids identified as upregulated (green bars) or downregulated (red bars) in activated ChAT^+^ mice (F) Annotations of bile acids highlighted in (E). Metabolite IDs are colored according to annotation as primary (blue) or secondary (orange) bile acids. Metabolite IDs are specific to each sample (N=12-14 for each group analyzed, *: p<0.05, **: p<0.01, ***: p<0.001, ****: p<0.0001) See also Table S1

To contextualize these data, we applied the Global Natural Products Social Molecular Networking (GNPS) tool^47^, an open-access mass spectrometry repository and analysis pipeline. GNPS revealed metabolic networks of both annotated and unannotated molecules in the cecal contents of ChAT^+^-activated and TH^+^-activated mice (Figures 3C and 3D), allowing us to identify metabolites with differential abundance between control and activated samples. Activation of TH^+^ neurons strongly increased metabolites whose closest spectral matches were linoelaidic acid (ID: 626), oleanolic acid methyl ester (ID: 378), and coproporphyrin I (ID: 739). Metabolites that spectrally resembled xanthine (ID: 259), genistein (ID: 846), and trans-ferulic acid (ID: 707) were decreased upon activation of TH^+^ neurons (Table S1).

In both ChAT^+^-activated and TH^+^-activated mice, the molecular networks largely consisted of level 3 annotations (based on the Metabolomics Standards Initiative (MSI)^48^) of compounds belonging to the bile acid molecular family and their conjugates, as well as unannotated analogs (Figures 3C-3D). Primary bile acids are chemicals derived from host (mouse) cholesterol biosynthesis, which are subsequently co-metabolized by gut bacteria into secondary bile acids^49,50^. Interestingly, metabolites with a closest spectral match to the primary bile acid cholic acid (IDs: 108, 114, 215, 219, 221, 224, 259) were significantly enriched in the cecum of ChAT^+^-activated mice (Figures 3D-3F; Table S1). Additional metabolites that spectrally resemble tauro-conjugated primary bile acids, such as taurocholic acid (IDs: 234, 248) and taurohyocholic acid (ID: 235), trended upwards. Conversely, features matching the spectra of secondary bile acids and bile acid metabolites such as ursodeoxycholic acid (ID: 13), deoxycholic acid (ID: 100), beta-hyodeoxycholic acid (IDs: 1, 143) and 12-ketodeoxycholic acid (IDs: 19, 138) were decreased in ChAT^+^-activated mice (Figures 3D-3F). These data suggest that activation of ChAT^+^ neurons may modulate, either directly or indirectly, primary bile acid secretion and/or metabolism to secondary bile acids, which have been implicated in a number of metabolic and immunologic functions, as discussed below.

### Neuronal subpopulations differentially shape the gut luminal proteome

Proteins from the mouse, gut microbes, and diet converge and interact in the GI tract^51^. We performed untargeted label-free proteomics by LC-MS/MS of cell-free supernatants of the cecal contents from ChAT^+^-activated and TH^+^-activated mice and controls collected one hour following the final C21 treatment (see Figure 2A). Consistent with the increase in cecal bile acid metabolites that we observed in ChAT^+^-activated mice, we report an increased abundance of Niemann-Pick C1-Like 1 protein (NPC1L1) in the cecum of these mice (Figure 4A). NPC1L1 is expressed on the apical surface of enterocytes, and is integral to the absorption of free cholesterol, the precursor of bile acids, from the lumen^52^. Goblet cell-related proteins, specifically Mucin-19 (MUC19) and Zymogen granule 16 (ZG16), a protein localized to secretory granules^53^, also trended upwards following ChAT^+^ neuronal activation (Figure 4A). Conversely, one of the most highly downregulated proteins was an aldehyde dehydrogenase (Q3U367) encoded by the *Aldh9a1* gene, which is involved in the catalytic conversion of putrescine to gamma-aminobutyric acid (GABA)^54^. While GABA is the primary inhibitory neurotransmitter in the CNS, little is known about its role in the ENS. The most significantly upregulated proteins in cecal contents of ChAT^+^-activated mice were pancreatic digestive enzymes including chymopasin (CTRL), chymotrypsinogen B1 (CTRB1), and pancreatic lipase related protein 2 (PNLIPRP2) (Figure 4A). Accordingly, network analysis of upregulated proteins revealed that KEGG pathways associated with digestion represent the majority of the network (Figure 4B). This is consistent with evidence that cholinergic, viscerofugal neurons send signals from the GI tract to other organs of the digestive system, including the pancreas^4^. Cholinergic innervation of the pancreas plays a significant role in regulating pancreatic functions, such as the secretion of digestive enzymes and insulin release^55^.

**Figure 4.**
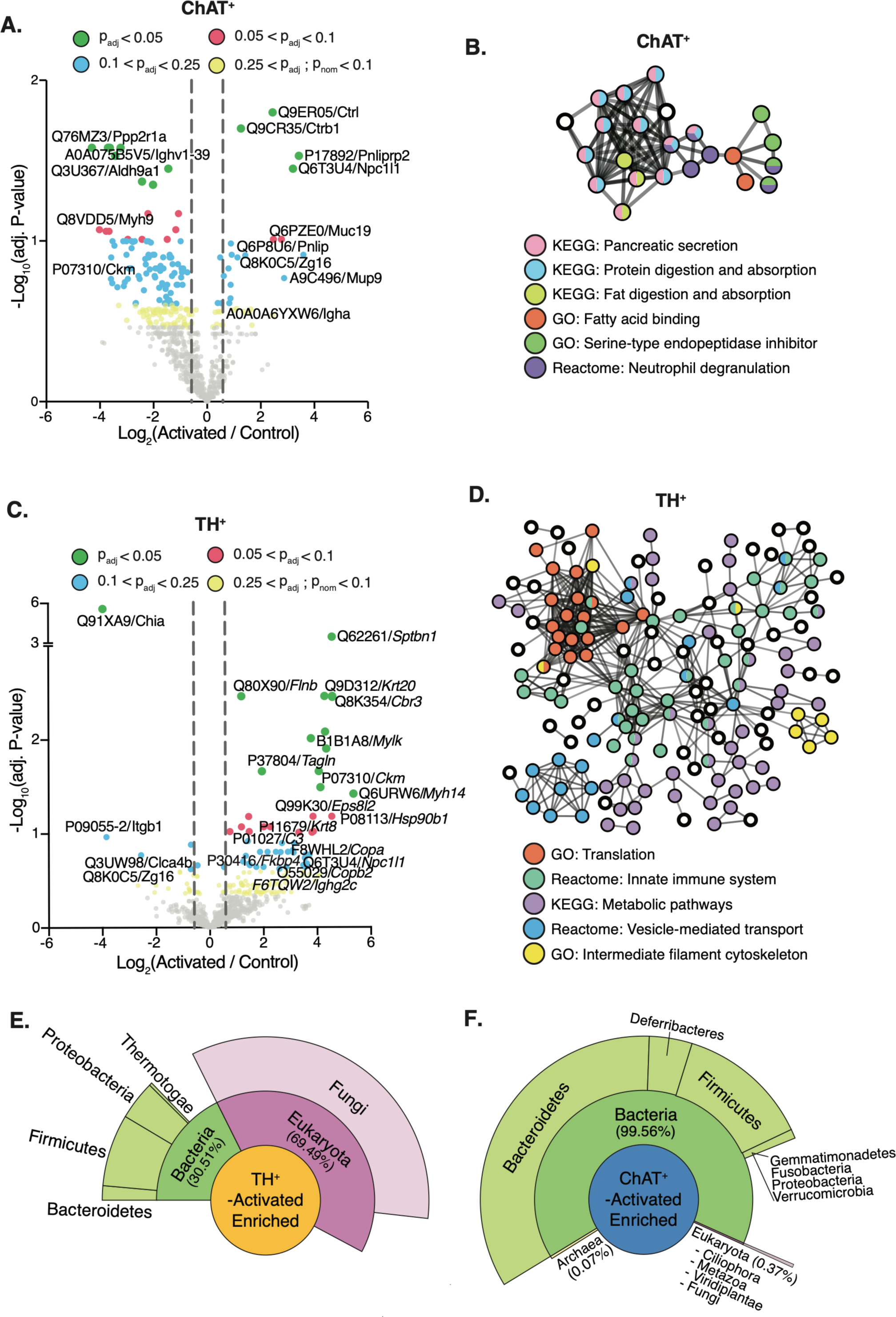
Gut-Associated ChAT^+^ or TH^+^ Neuronal Activation Alters Host and Microbe-Derived Luminal Proteins. (A) Volcano plot of differentially-expressed host proteins identified in the cecal contents of ChAT^+^-activated (N=8) vs. ChAT^+^-control mice (N=9) mice, 1 hour after final C21 administration (B) STRING network analysis of host proteins that were more abundant in ChAT+-activated mice (p_nom._< 0.2) (C) Proteomic volcano plot of TH^+^-activated vs. TH^+^-control mice (N=7 mice per group) (D) STRING network analysis of upregulated host proteins in TH^+^-activated mice (p_nom._< 0.2). (E-F) Unipept metaproteomic analysis of upregulated microbial proteins (fold change > 2, p_nom._< 0.2) in TH^+^-activated and ChAT^+^-activated mice (N=7-9 mice per group) **Source Data** Figure 4A https://github.com/mazmanianlab/Griffiths_Yoo_et_al/blob/main/proteomics/CHAT_proteomics_volcano.txt **Source Data** Figure 4C https://github.com/mazmanianlab/Griffiths_Yoo_et_al/blob/main/proteomics/TH_proteomics_volcano.txt **Source Data** Figures 4E **and 4F** https://github.com/mazmanianlab/Griffiths_Yoo_et_al/blob/main/proteomics/metaproteomics/Microbiome_associated_proteins.xlsx

Peripheral activation of TH^+^ gut-associated neurons also altered the luminal proteome of the cecum. Notably, 88% (52/59) of the differentially abundant proteins (p_adj._<0.25) were distinct from those identified in ChAT^+^-activated mice. The overall direction of the effect was also reversed: ∼90% of differentially-abundant cecal proteins in TH^+^-activated mice were upregulated (53/59), compared to ∼18% in ChAT^+^-activated mice (20/112), suggesting that activation of distinct neuronal subsets is associated with opposing changes in GI function. We observed signatures of increased protein-protein interactions in cecal contents of TH^+^-activated mice, evidenced by more network nodes and connections (Figure 4D). Filamin B (FLNB) and spectrin beta chain, non-erythrocytic 1 (SPTBN1) were two of the most significantly enriched proteins following TH^+^ neuron activation (Figure 4C). Both are associated with the intestinal brush border and membrane vesicles^56,57^. Accordingly, coatomer proteins also trended upward (COPA and COPB2) (Figure 4C) and vesicle-mediated transport was one of the major protein networks altered (Figure 4D). Other upregulated protein interaction networks were associated with metabolic pathways, ribosomal activity, and the immune system (Figure 4D). For example, the immune-related proteins immunoglobulin heavy constant alpha (IGHA) (in ChAT^+^-activated), immunoglobulin heavy constant gamma 2C (IGHG2C), and complement component 3 (in TH^+^-activated) trended upward (Figures 4A and 4C).

Perhaps the most intriguing observation was the strong depletion of acidic mammalian chitinase (CHIA) upon activation of TH^+^ neurons (Figure 4C). Chitin is a natural polysaccharide that is a major component of fungal cell walls^58^, but intestinal chitinases are poorly studied in mice. This result prompted us to query the pan-proteomic dataset against a microbial protein database, which revealed that the decrease in CHIA abundance following TH^+^ neuron activation was accompanied by a large bloom in fungal-associated peptides in the microbiome (∼59% of peptides mapped to any microbe) (Figure 4E). In contrast, fungal peptides represented only ∼0.4% of enriched peptides in the lumen of ChAT^+^-activated mice (Figure 4F). Unfortunately, we were unable to corroborate these proteomic data with metagenomics since the DNA extraction method we used was not optimized for fungi. However, these findings suggest that the reduced chitinase production of activated TH^+^ cells is directly associated with a dramatic increase in fungal proteins, which, if experimentally validated in future, would represent a circuit by which the gut-associated neurons of mice regulates fungal load in the gut.

### Activation of ChAT^+^ and TH^+^ neurons alters the intestinal transcriptome

Given the changes to the gut microbiome, proteome and metabolome that we observed, we were interested in the tissue-level impact of neuronal activation on the intestinal transcriptome. We therefore profiled gene expression with QuantSeq, a quantitative 3’ mRNA-sequencing technology, in 1 cm of tissue from the distal SI and proximal colon harvested one hour after the last C21 injection. Rapid and transient expression of immediate early genes (IEGs) is widely used as a measure of increased neuronal activity^59^, and the IEGs *Fos*, *Egr1*, *Jun*, and *Klf2* were among the most significantly upregulated transcripts we identified in the SI and colon of both ChAT^+^- and TH^+^-activated mice (Figures 5A-5D). These IEGs are also known to be upregulated during growth and differentiation of highly active cell types such as immune cells^60,61^, smooth muscle cells^62^, and intestinal epithelial cells^63^.

**Figure 5.**
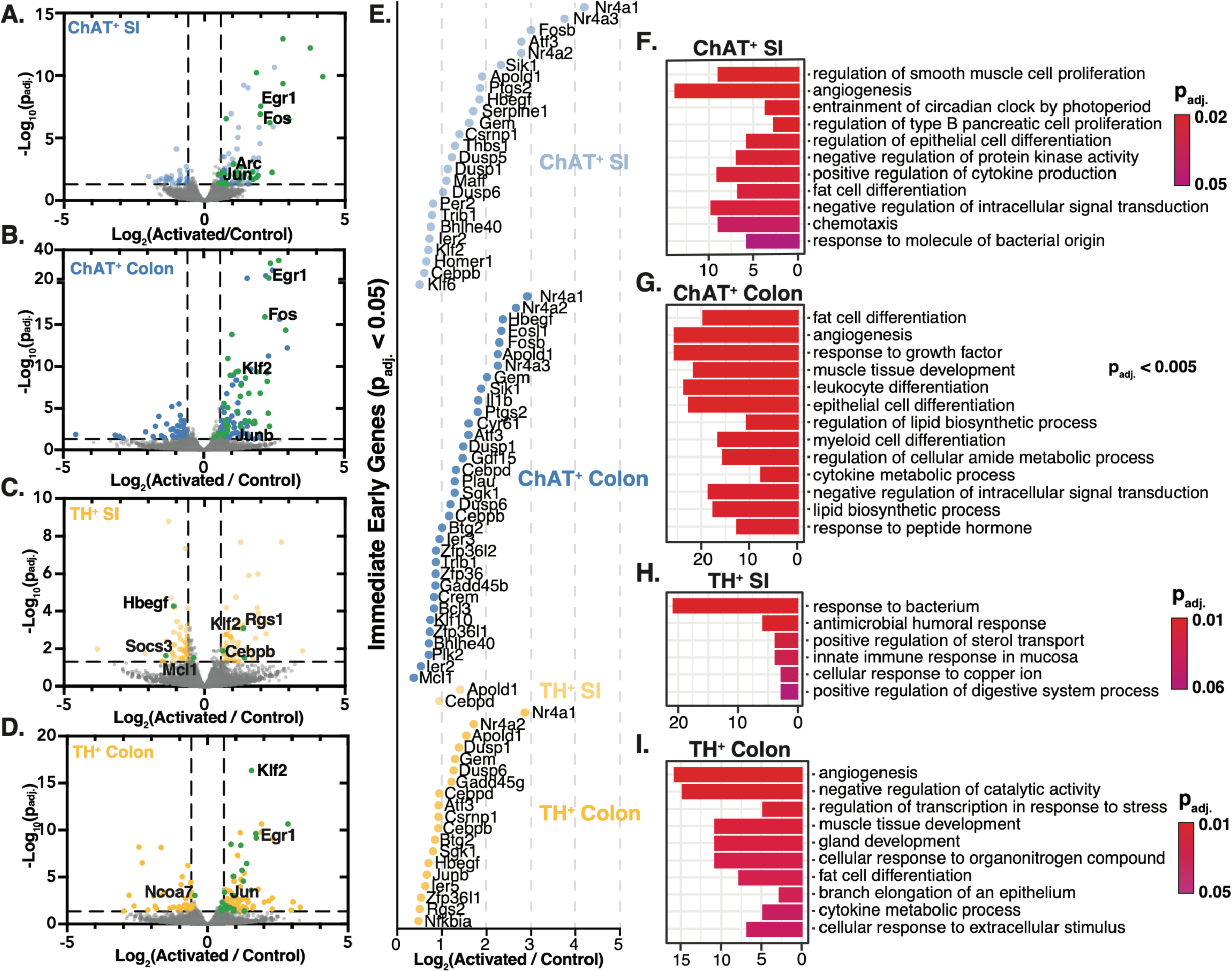
ChAT ^+^ and TH^+^ Activation-Mediated Transcriptomic Changes. (A-D) Differentially-expressed genes in DREADD-activated vs. control (A) ChAT ^+^ distal SI, (B) ChAT^+^ proximal colon, (C) TH^+^ distal SI, (D) TH^+^ proximal colon (Dashed vertical lines: Fold Change (FC) = +/-1.5; dashed horizontal lines: p_adj._< 0.05. Transcripts of IEGs are highlighted in green and annotated. N=10 mice per group) (E) Fold changes of upregulated IEGs (p_adj._< 0.05) as defined by ^59^ (F-I) Gene set enrichment analysis of gene ontology (GO) terms for (F) ChAT^+^ distal SI, (G) ChAT^+^ proximal colon, (H) TH^+^ distal SI, (I) TH^+^ proximal colon. See also Table S2 and Table S3 **Source Data** Figure 5 https://github.com/mazmanianlab/Griffiths_Yoo_et_al/tree/main/RNAseq

In the distal SI, we found similar numbers of differentially-expressed genes (DEGs; p_adj._<0.05) in ChAT^+^-activated mice (162 DEGs) and TH^+^-activated mice (165 DEGs) (Figures 5A and 5C). The direction of regulation differed, however, with ∼73% of DEGs upregulated upon ChAT^+^ activation (118 up, 44 down) and ∼58% of DEGs downregulated upon TH^+^ activation (69 up, 96 down). IEGs followed this overall pattern, with 29 upregulated in the distal SI of ChAT^+^-activated mice but only two upregulated in TH^+^-activated mice (Figure 5E), and three (i.e., *Hbegf, Soca3, Mcl1*) downregulated (Figure 5C). Similar proportions of DEGs were upregulated in the proximal colon of both ChAT^+^-activated (169 up, 84 down) and TH^+^-activated mice (130 up, 62 down) (Figures 5B and 5D). Given the enrichment in fungal proteins and reduction in the level of the CHIA protein in the TH^+^-activated mice, we explored potential immune responses to fungi but found no obvious inflammatory signals compared to control mice (Table S3).

To gain insight into the cellular functions of DEGs, we used Gene Set Enrichment Analysis (GSEA) (Figures 5F-I; Table S2). Notably, the most highly enriched gene ontology (GO) term for the distal SI of ChAT^+^-activated mice was “regulation of smooth muscle cell proliferation” (Figure 5F), whereas in TH^+^-activated mice it was “response to bacteria” (Figure 5H), consistent with the increase in immune-related responses suggested by our proteomic dataset. In the proximal colon, we observed similar GO pathways in ChAT^+^-activated and TH^+^-activated mice (Figures 5G and 5I), suggesting that transcriptomic signatures may depend on the context of the activated neurons. In the SI, ChAT^+^ neurons predominantly border muscle cells in the myenteric plexus, while TH^+^ neurons neighbor epithelial and immune cells which respond to bacteria in the submucosal plexus of the distal SI (see Figures 1C-1E). In the colon, both neuronal subsets are abundant in the myenteric plexus (see Figure 1D). In both myenteric and submucosal plexuses, we saw a wider breadth of pathways upregulated by activation of ChAT^+^ neurons than TH^+^ neurons, with ChAT^+^ neuronal activation impacting diverse cellular functions in the GI tract, involving endothelial, epithelial, immune, and adipose cells (Figure 5G; Table S2).

### Differential functional GI outcomes of activation of ChAT^+^ and TH^+^ neurons

Motivated by the complexity of responses we observed following activation of neuronal populations in the gut, we decided to assay functional GI outcomes. Both ChAT^+^ and TH^+^ neuronal populations are known to be important for motility and secretory function^29,64^, but they have never been specifically modulated to study GI physiology in a freely behaving mammal. Activation of either ChAT^+^ or TH^+^ gut-associated neurons resulted in faster whole gut transit time, increased fecal pellet output, and mass of cecal contents compared to control mice (Figures 6A-C). Fecal pellets from ChAT^+^-activated, but not TH^+^-activated, mice had increased water content, which is consistent with reports in the literature of involvement of ChAT^+^ enteric neurons in fluid secretion (Figures 6D-6F)^15,23,24^. This distinction is particularly notable given the higher concentration of TH^+^ neurons than ChAT^+^ neurons in most regions of the submucosal plexus (see Figure 1), which is largely responsible for fluid secretion and absorption^1^. Daily administration of C21 for 9 days to control mice (no DREADD) did not cause any obvious health impairment and the mice maintained body weight throughout the experimental period (Figures S6C and S6D). TH^+^-activated mice also maintained body weight, but ChAT^+^-activated animals experienced slight weight loss that likely reflects the diarrhea-like phenotype over 9 consecutive days. To further examine gut motility in the absence of extrinsic innervation, we analyzed propulsive colonic migrating motor complexes (CMMCs) in an *ex vivo* system. Activation of ChAT^+^ neurons by C21 administration resulted in more frequent migration of motor complexes (Figures 6G, 6I, and 6J; Table S4), whereas activation of TH^+^ neurons had no effect on CMMCs (Figure 6H; Table S4). The discrepancy between the *in vivo* and *ex vivo* results we observed may be due to activation of TH^+^ neurons in the sympathetic prevertebral ganglia that project to the gut^32^. Overall, these data reveal that ChAT^+^, but not TH^+^, neurons in the gut mediate intestinal fluid balance and *ex vivo* colonic motility.

**Figure 6.**
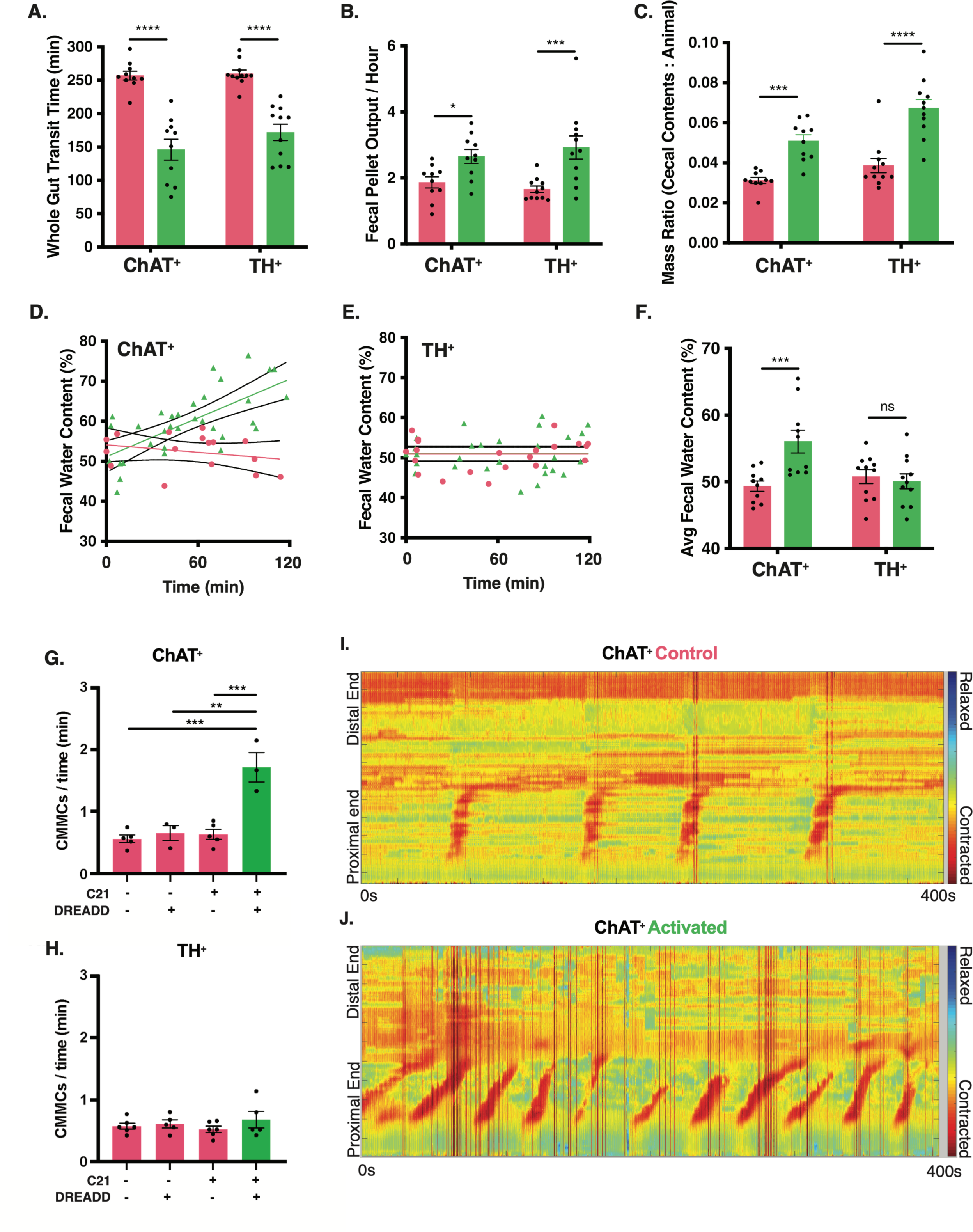
GI Physiology Differences in ChAT^+^ vs. TH^+^ mice Following Activation. (A) Activation-mediated changes in whole gut transit time in ChAT^+^ and TH^+^ mice (B) Activation-mediated changes in fecal pellet output in ChAT^+^ and TH^+^ mice (C) Activation-mediated changes in normalized cecal content mass in ChAT^+^ and TH^+^ mice (D-E) Fecal pellet water content in (D) ChAT^+^ and (E) TH^+^ mice over 2 hours following C21 activation with a least squares nonlinear regression displaying 95% confidence interval (F) Average fecal pellet water content in ChAT^+^ and TH^+^ mice following activation (A-F: N = 10-11 mice per group; *: p<0.05, ***: p<0.001, ****: p<0.0001, determined by 2-way ANOVA with Sidak’s method for multiple comparisons) (G-H) Frequency of *ex vivo* CMMCs from (G) ChAT^+^ and (H) TH^+^ mice over 30 minutes following activation (N = 3-6 mice per group; **: p<0.01, ***: p<0.001, determined by 2-way ANOVA with Sidak’s method for multiple comparisons) (I-J) Heatmaps showing frequency of CMMCs over 400 seconds following activation in *ex vivo* preps from (I) ChAT^+^ control and (J) ChAT^+^ DREADD-administered mice See also Figure S6 and Table S4

## DISCUSSION

While early pioneers of neuroscience in the 20^th^ century focused on the ENS as a model, more recent research has centered on the brain, and our understanding of the CNS has outpaced that of other neuronal systems in the body. As a result, basic knowledge of many aspects of neuronal architecture and function within the gut remain rudimentary^20,65,66^. Here, using a viral delivery system with enhanced tropism for the ENS, we mapped the distribution of ChAT^+^ and TH^+^ neurons across the mouse GI tract and assayed the complex effects of their peripheral activation on physiology and function. Although the DREADD-based activation paradigm we use in this study is inherently artificial, the results reveal strikingly different roles for neuronal populations, with nearly every feature characterized (spatial distribution, metagenomic, metabolic, transcriptional, and proteomic profiles, and even physiological output) unique to cell type.

The viral vector we used, AAV-PHP.S, can transduce other neuronal subtypes in the PNS such as those in the DRGs and, with a strong ubiquitous promoter, induce transgene expression^30^. To limit this off-target effect, we utilized a weaker promoter with increased ENS specificity and focused our analyses on GI tissue and lumen. By thus excluding most known extrinsic innervation pathways, we uncover cell-type-specific effects of gut-associated neuronal activation that are independent of signaling from the brain. Exposure to the external environment charges the intestines with myriad responsibilities including absorption and digestion of dietary nutrients, exclusion of xenobiotics, protection from enteric infection, and partnership with the gut microbiome. Deletion of ChAT in enteric neurons leads to microbiome dysbiosis^67^, and we observed differences in the compositional profile (both metagenomic and proteomic) of the gut microbiome specific to activation of ChAT^+^ neurons. Notably, we found an expansion of Verrucomicrobia driven by *A. muciniphila*, which has been implicated in human diseases such as obesity^68,69^, multiple sclerosis^70,71^, and seizures^72^. *A. muciniphila* metabolizes host-derived mucus as a nutrient source^1^, consistent with the increase in luminal mucin proteins and digestive enzymes we observed in ChAT^+^-activated mice. A particularly interesting host-microbial interaction emerged from activation of TH^+^ cells, which led to a dramatic decrease in anti-fungal chitinase (CHIA) protein expression and a concomitant bloom in fungi, suggesting that neuronal circuits can regulate fungal populations in the gut. If validated, this intriguing host-microbial interaction could have implications for health. Our study does not, however, reveal the mechanism(s) by which ENS activation reshapes the gut microbial community structure, which may involve altered colonic motility, changes in mucus production, modulation of mucosal immune responses, and/or shifts in metabolism and nutrient availability.

The human gut microbiome possesses as much metabolic capacity as the liver; it is therefore no surprise that changes to both mouse gut physiology and the microbiome have major influences on the gut metabolome. In a striking example of mutualism, we report widespread changes to the pool of intestinal bile acids, molecules produced via host-microbial co-metabolism. Activation of ChAT^+^ neurons, but not TH^+^ neurons, impacted expression of NPC1L1, which is involved in cholesterol transport. In mammals, cholesterol is the substrate for production of primary bile acids, which are then metabolized exclusively by the gut microbiome into secondary bile acids. Bile acids play critical roles in fat absorption^73^, gut motility^74^, hormonal signaling^75^ and immune functions^76^, and neurological conditions^77^. Expression of bile salt hydrolases and hydratases increases the fitness of both commensal and pathogenic bacteria^78–82^. While additional work is required to determine how the ENS affects levels and constitution of the bile acid pool, understanding the processes that regulate synthesis of secondary bile acids may have implications for organ systems throughout the body.

Our study complements recent single-cell RNAseq studies of the ENS neuronal transcriptome^83^ by giving us the ability to selectively activate specific enteric neurons and explore the dynamic interplay between cells of various lineages in the gut. Importantly, the transcriptomic changes we observed may be a consequence of direct or indirect effects of neuronal activation. Indeed, induced activation of ChAT^+^ or TH^+^ neurons rapidly changed GI transit and fluid secretion patterns, which are only a fraction of the processes that may feed back on epithelial or immune cells, altering their gene expression profiles. Further single-cell analysis may help dissect the roles of the various intestinal cells that collaborate to coordinate gut functions.

The ENS adapts and responds to incredibly diverse molecular cues from the environment and must do so throughout the entire length and surface area of the intestines—the largest and most extensive internal organ, with a rich network of neurons termed the “second brain” (Gershon, 2015). Exposure to molecules from the diet or the microbiome may modulate ENS function, along with signals from outside the gut such as the circulatory system. Curiously, many disorders of the brain are also associated with GI symptoms^84–88^. While mechanisms linking the gut and the brain, and their consequences for health, are an active area of study, the impact of neuronal activation within the ENS has largely been unexplored. Herein, we establish an experimental system that allows controlled activation of intrinsic and extrinsic neurons of the gut, separated from inputs from the brain, and demonstrate broad changes in the gut environment and its physiology that differ by activated neuronal population. The extensive datasets on activation of two major gut-associated neuronal populations that we generated should serve as a resource for further studies on the interconnected biological systems governing the complex relationship between gut physiology and the microbiome. Future deployment of this approach could enable mapping of neuronal connections into and out of the gut, providing insights into how the ENS networks with tissues throughout the body and advancing growing research into the many functions of the GI tract, an endeavor with important consequences for human health.

### Limitations of the Study

It was recently shown that a similar strategy using AAV-PHP.S-hSYN1 in ChAT-Cre mice also labels ChAT^+^ neurons in cardiac ganglia, and activation of cholinergic neurons reduces heart rate and blood pressure^89,90^. Cardiac afferent neurons signal through the vagus nerve and jugular-nodose ganglia to the brain, and in sensory pathways through the DRGs and spinal cord^91^. Though there is no known direct route for signaling from cardiac ganglia to gut-associated neurons, it is possible that the gut may be impacted by an indirect route involving the CNS. In the future, further refinement of AAVs through directed evolution may generate serotypes with exclusive tropism for the ENS and allow full separation of the functions of intrinsic and extrinsic activation of gut-associated neuronal subsets. Since the hSYN1 promoter we used has been shown to drive expression only in neurons^92,93^, we did not characterize other cell types, such as enteroendocrine cells (EECs), also called neuropod cells, which were recently shown to form synapses with enteric neurons and contribute to sensory transmission from the gut to the brain through the vagus nerve. Since these cells turn over every few days^94^ and have not been reported to express TH or ChAT, we think it unlikely that they are a major contributor to the activation-induced phenotypes we observed, but we cannot completely rule out their involvement.

## ACKNOWLEDGMENTS

We thank members of the Mazmanian laboratory and Dr. Jonathan Hoang for discussions throughout the research project and critical reading of the manuscript, and Dr. Catherine Oikonomou for invaluable manuscript editing. We thank Dr. Andres Collazo and Caltech’s Biological Imaging Facility for training and access to microscopy capabilities and the Caltech Proteome Exploration Laboratory for access to LC-MS/MS capabilities. We thank Professor Elisa Hill-Yardin for providing MATLAB scripts for CMMC heatmaps. This research was funded in part by Aligning Science Across Parkinson’s (ASAP-020495 and ASAP-000375) through the Michael J. Fox Foundation for Parkinson’s Research (MJFF). For the purpose of open access, the authors have applied a CC BY 4.0 public copyright license to all Author Accepted Manuscripts arising from this submission. S.K.M. was supported by grants from the Heritage Medical Research Institute, Emerald Foundation, Caltech Center for Environmental and Microbial Interactions (CEMI), the National Institutes of Health (GM007616 and DK078938), and the Department of Defense (PD160030). P.C.D. was supported by grants from National Institute of Diabetes and Digestive and Kidney Diseases (R01DK136117 and U24DK133658).

## AUTHOR CONTRIBUTIONS

J.A.G., B.B.Y., and S.K.M. designed experiments. J.A.G. and B.B.Y. performed experiments. K.Y.C., C.C. (supervised by V.G.), and J.A.G. helped with AAV-mediated ENS characterization. P.T.B (supervised by D.W.W.), M.J.S., and A.M. performed acquisition and analysis of proteomic data. V.C., G.H., G.S., Q.Z., and J.S. (supervised by R.K.) performed acquisition and analysis of metagenomic data. K.W. (supervised by P.C.D.) performed acquisition and analysis of metabolomic data. T.M.T. assisted in animal-related work. J.A.G., B.B.Y., and S.K.M wrote the manuscript with input from all authors.

## METHODS

### Resource availability

Lead contact Further information and requests for resources and reagents should be directed to and will be fulfilled by the lead contact, Sarkis K. Mazmanian.

### Materials availability

No new reagents were generated in this study.

### Data & Code Availability

Microbial sequencing data have been deposited at the European Bioinformatics Institute (ERP131523), Metabolomic data at UCSD MassIVE repository (MSV000084550), proteomic data at UCSD MassIVE repository (MSV000087917), and QuantSeq data at NCBI GEO repository (GSE180961) and are publicly available as of the date of publication. All other experimental data used to generate the figures reported in this paper can be found at (https://github.com/mazmanianlab/Griffiths_Yoo_et_al/) and are publicly available as of the date of publication.

This paper does not report original code.

Any additional information required to reanalyze the data reported in this paper is available from the lead contact upon request.

### Experimental model and study participant details Mice

All mouse experiments were performed in accordance with the NIH Guide for the Care and Use of Laboratory Animals using protocols approved by the Institutional Animal Care and Use Committee at the California Institute of Technology. Mice were fed ad libitum for the entire duration of experiments. Homozygous TH-Cre (gift to V.G. from Ted Ebendal, B6.129X1-Thtm1(cre)Te/Kieg^96^) and ChAT-Cre (Jackson Laboratories, Bar Harbor, ME-Stock# 028861, RRID:IMSR_JAX:028861) mice were bred to wild-type mice to yield the male and female heterozygous Cre-mice used for our studies. Wild-type specific pathogen free (SPF) C57BL/6 (Jackson Laboratories, Bar Harbor, ME-Stock #000664, RRID:IMSR_JAX:000664) males and females were used for breeding and experiments. Mice at 6-8 weeks of age were used for experiments.

### Virus Production

Virus was produced using the methods described in Challis et al., 2019^42^ and dx.doi.org<u>/10.17504/protocols.io.bzn6p5he</u>. Briefly, human embryonic kidney (HEK293T) cells were triple-transfected with pUCmini-iCAP-AAV-PHP.S, pHelper plasmid, and one of the following pAAV genomes: hSYN1-tdTomato, hSYN1-mRuby2, hSYN1-DIO-mRuby2, hSYN1-mNeonGreen, CAG-mNeonGreen, hSYN1-DIO-mNeonGreen, hSYN1-mTurquoise2, hSYN1-DIO-mTurquoise2, hSYN1-DIO-hM3Dq-mRuby2, CAG-GCaMP6f. Cells were grown in DMEM + Glutamax + Pyruvate (Gibco, Gaithersburg, MD-Stock# 10569-010) + 5% FBS + non-essential amino acids (Gibco, Gaithersburg, MD-Stock# 11140-050) + penicillin-streptomycin (Gibco, Gaithersburg, MD-Stock# 15070-063). Virus was precipitated from cells and supernatant with an 8% PEG solution (wt/vol), and purified by ultracentrifugation using 15%, 25%, 40%, 60% stacked iodixanol gradients.

### Method Details

#### Systemic Delivery of AAV

Mice were anesthetized using 2% isoflurane. Virus was titered to 1×10^12^ vg, resuspended in a volume of 100 µl with sterile PBS, and injected retro-orbitally as described in dx.doi.org/<u>10.17504/protocols.io.bzn6p5he</u>.

### Neuronal activation of the GI tract

See dx.doi.org/<u>10.17504/protocols.io.bzp5p5q6</u>. TH-Cre and ChAT-Cre mice were used for these experiments. “Activated” mice were infected with AAV-PHP.S-hSYN1-DIO-hM3Dq-mRuby2 and “Control Mice” were infected with AAV-PHP.S-hSYN1-DIO-mRuby2. This was to control for both AAV-PHP.S-mediated expression and the effects of Compound 21 dihydrochloride (C21) (HelloBio, Princeton, NJ-HB6124). C21 was injected intraperitoneally (i.p.) at a dose of 3 mg/kg for 10 consecutive days to both groups of mice. Mice for timecourse experiments were single-housed in sterile cages with autoclaved water following the first C21 administration. Injections of C21 were administered at the same time every day (10AM). Mice were sacrificed one hour after the day 10 injection.

### Tissue Preparation, Immunohistochemistry, Imaging, and Quantification

Procedures are described in dx.doi.org/10.17504/protocols.io.bzp6p5re. 100 mg/kg of pentobarbital (Euthasol - Virbac, Carros, France) was administered i.p., and tissues were perfused with 30 mL of phosphate-buffered saline (PBS) and then cold 4% paraformaldehyde (PFA) in PBS. GI tract was post-fixed in 4% PFA overnight at 4 °C, and stored in PBS + 0.025% sodium azide. Tissues that underwent subsequent immunohistochemistry were made transparent by the passive CLARITY technique (PACT) as in Treweek et al., 2015^35^. Briefly, perfused and fixed tissues were embedded with polymerized 4% (wt/vol) acrylamide, and lipids were eliminated using 8% (wt/vol) SDS solution. Jugular-nodose ganglia and dorsal root ganglia tissues were cryoprotected in 10% then 30% sucrose in PBS for 1 day each. Tissues were embedded and flash frozen in OCT and cryostat sectioned into 40 µM sections. Spinal cord and brain tissues were vibratome sectioned into 50 µM sections. Tissues were blocked in 3% donkey serum and permeabilized with PBS + 0.3% Triton (PBST). Primary antibodies were incubated in PBST for 48 hours and washed with PBST for 24 hours (replacing the wash solution 3 times). Tissues were next incubated in secondary antibodies (and DAPI) for 24 hours and washed in PBS for 48 hours, intermittently replacing the wash solution with fresh PBS. Primary antibodies used were rabbit anti-PGP9.5 (1:300; Millipore Cat# AB1761-I, RRID:AB_2868444), rabbit anti-tyrosine hydroxylase (1:500, Abcam Cat# ab112, RRID:AB_297840), rabbit anti-choline acetyltransferase, (1:250, Abcam Cat# ab178850, RRID:AB_2721842), and mouse anti-NeuN (1:300, Abcam Cat# ab104224, RRID:AB_10711040). Secondary antibodies used were donkey anti-rabbit Alexa 568 (Thermo Fisher Scientific Cat# A10042, RRID:AB_2534017), goat anti-rabbit Alexa 647 (Thermo Fisher Scientific Cat# A-21245 (also A21245), RRID:AB_2535813), and goat anti-mouse Alexa 594 (Thermo Fisher Scientific Cat# A-11032, RRID:AB_2534091). GI tissues imaged for virally-expressed, endogenous fluorescence were made transparent using a sorbitol-based optical clearing method, Sca*l*eS as in Hama et al., 2015^34^. Tissues were mounted in method-respective mounting media (RIMS and Scales S4) on a glass slide with a 0.5mm spacer (iSpacer, SunJin Lab Co.). Images were acquired on Zeiss LSM 780 or 880, and microscope, laser settings, contrast, and gamma remained constant across images that were directly compared. All confocal images were taken with the following objectives: Fluar 5× 0.25 M27 Plan-Apochromat, 10× 0.45 M27 (working distance 2.0 mm) and Plan-Apochromat 25× 0.8 Imm Corr DIC M27 multi-immersion. Neurons in each ganglion were counted by counting cells that were of a distinct color. Colonic ganglia were defined as distinct if separated by a width of 3 or more neurons.

### GCaMP6f Fluorescence in *Ex Vivo* Intestinal Preparations

As described in dx.doi.org/<u>10.17504/protocols.io.bzqap5se</u>, small intestinal tissue was quickly harvested from ChAT-Cre mice, flushed, and placed in oxygenated (95% O_2_, 5% CO_2_), ice cold Krebs-Henseleit solution for 1 hour followed by 15 min at room temperature. A segment was cut along the mesenteric attachment and pinned flat (mucosa facing down) on a Sylgard-lined recording chamber (Warner Instruments, PH1) in oxygenated Krebs-Henseleit solution. C21 was added at 10nM and GCaMP6f fluorescence was detected on an upright microscope (Zeiss, Oberkochen, Germany-Examiner D1).

### Metagenomics

Procedures are described in dx.doi.org/10.17504/protocols.io.bzqep5te.

#### Fecal collection

AAV-PHP.S-hSYN1-DIO-hM3Dq-mRuby2 (10^12^ vg) was delivered systemically to TH-Cre and ChAT-Cre mice. 3-4 week after infection, C21 (3 mg/kg) was administered daily for 10 consecutive days. Fecal pellets were collected in sterile containers one day before the initial C21 dose, and on day 2, 6, and 10 of injections.

#### Fecal sample DNA extraction and library preparation

DNA was extracted with the Qiagen MagAttract PowerSoil DNA kit as previously described in Marotz et al., 2017^118^.This protocol is optimized for an input quantity of 1 ng DNA per reaction. Prior to library preparation, input DNA was transferred to a 384-well plate and quantified using a PicoGreen fluorescence assay (ThermoFisher, Inc). Input DNA was then normalized to 1LJng in a volume of 3.5 µL of molecular-grade water using an Echo 550 acoustic liquid-handling robot (Labcyte, Inc). Enzyme mixes for fragmentation, end repair and A-tailing, ligation, and PCR were prepared and added in approximately 1:8 scale volumes using a Mosquito HV micropipetting robot (TTP Labtech). Fragmentation was performed at 37 °C for 20LJmin, followed by end repair and A-tailing at 65 °C for 30LJmin.

Sequencing adapters and barcode indices were added in two steps, following the iTru adapter protocol^119^. Universal “stub” adapter molecules and ligase mix were first added to the end-repaired DNA using the Mosquito HV robot and ligation performed at 20 °C for 1 h. Unligated adapters and adapter dimers were then removed using AMPure XP magnetic beads and a BlueCat purification robot (BlueCat Bio). 7.5 µL magnetic bead solution was added to the total adapter-ligated sample volume, washed twice with 70% EtOH, and then resuspended in 7 µL molecular-grade water.

Next, individual i7 and i5 were added to the adapter-ligated samples using the Echo 550 robot. Because this liquid handler individually addresses wells, and we used the full set of 384 unique error-correcting i7 and i5 indices, we were able to generate each plate of 384 libraries without repeating any barcodes, eliminating the problem of sequence misassignment due to barcode swapping^120,121^. To ensure that libraries generated on different plates could be pooled if necessary, and to safeguard against the possibility of contamination due to sample carryover between runs, we also iterated the assignment of i7 to i5 indices each run, such that each unique i7:i5 index combination was repeated only once every 147,456 libraries. 4.5 µL of eluted bead-washed ligated samples was added to 5.5 µL of PCR master mix and PCR-amplified for 15LJcycles. The amplified and indexed libraries were then purified again using magnetic beads and the BlueCat robot, resuspended in 10 µL water, and 9 µL of final purified library transferred to a 384-well plate using the Mosquito HV liquid-handling robot for library quantitation, sequencing, and storage. 384 samples were then normalized based on a PicoGreen fluorescence assay.

#### Shallow shotgun metagenome sequencing and diversity analysis

The Illumina data for each HiSeq lane was uploaded to Qiita, a tool with standardized pipelines for processing and analyzing metagenomic data^122^. Adapter sequences were removed from the reads using the Atropos v.1.1.15 (RRID:SCR_023962, https://github.com/jdidion/atropos)^123^ command (from the qp-shogun 0.1.5 pipeline) and the trimmed sequences were downloaded from Qiita. The reads for each sample were filtered of any potential mouse contamination using Bowtie2 v.2-2.2.3^98^ (RRID:SCR_016368, https://bowtie-bio.sourceforge.net/bowtie2/index.shtml). The filtered reads were then aligned to the Web of Life (WoL) reference phylogeny^124^ with Bowtie2 using an adapted SHOGUN pipeline^125^. The WoL contains 10,575 bacterial and archaeal genomes, with each genome representing an operational taxonomic unit (OTU). Sequencing reads that did not map to a single reference genome as well as reads that mapped to multiple genomes were not included in the analysis. If an OTU had a relative abundance less than 0.01% in a given sample, the OTU was not included for that sample. Additionally, OTUs with fewer than 5 assigned reads were not considered. The samples were rarefied to a depth of 12,750 reads and those with fewer than the rarefaction depth were excluded. The QIIME2 v.2019.7^95^ (RRID:SCR_021258, https://qiime2.org/) DEICODE plugin was used to calculate the Aitchison distances, a compositional beta diversity metric, and perform Robust Aitchison PCA to create biplots that visualize relationships between features and samples^126^. The QIIME2 diversity plugin was used to calculate the other alpha- and beta-diversity metrics used in this study.

#### Metagenomics-based functional profiling

The filtered reads were also analyzed using HUMAnN2 v2.8.1^99^ (RRID:SCR_016280, https://huttenhower.sph.harvard.edu/humann2) to establish functional profiles for the samples. HUMAnN2 is a pipeline that begins by using MetaPhlAn2 to compile custom databases of reference genomes based on the species detected in a sample^127^. HUMAnN2 then maps the filtered reads onto these custom databases and the reads that do not map to any of the references are then subjected to a translated search against UniProt Reference Clusters or UniRef^100^ (RRID:SCR_010646, https://www.uniprot.org/help/uniref). Here, the UniRef90 database was used for the translated search and installed according to the HUMAnN2 documentation. The results from both the search performed using the custom reference genome database and the search against the UniRef90 database were combined and the gene families identified in each sample were reported in units of reads per kilobase (RPKs) to account for gene length. HUMAnN2 also compared the gene families found in a sample with the MetaCyc pathways database^128^ (RRID:SCR_007778, https://metacyc.org/) and output a table reporting the pathway abundances found in each sample. After rarefying gene family tables to a depth of 166,000 RPKs and using a depth of 22,600 for pathway abundances, the QIIME2 diversity and DEICODE plugins were used to calculate alpha- and beta-diversity metrics. The metagenomics data from this study are available from (https://github.com/mazmanianlab/Griffiths_Yoo_et_al/tree/main/metagenomics).

### Metabolomics

Procedures are described in dx.doi.org/10.17504/protocols.io.bzqfp5tn.

#### Sample preparation

Frozen cecal samples were transported on dry ice for metabolomics analysis. The samples were weighed and an extraction solvent (1:1 methanol to water with an internal standard of 1 µM sulfamethazine) was added at a 1:10 milligram to microliter ratio. The samples were then homogenized using a TissueLyser II (Qiagen) for 5 minutes at 25 Hz followed by a 15 min centrifugation at 14,000 rpm. 120 µL of supernatant was transferred to a 96-well DeepWell plate (Eppendorf) and lyophilized using a CentriVap Benchtop Vacuum Concentrator (Labconco) and stored at −80 °C. At the time of data acquisition, the lyophilized plates were resuspended in a 1:1 methanol to water solvent spiked with 1 µM of sulfadimethoxine. The plates were vortexed for 2 minutes, centrifuged at 14,000 rpm for 15 min and 120 µL of the supernatant was transferred to a 96-well autosampler plate (Eppendorf). Plates were stored at 4 °C prior to LCMS analysis.

#### Data acquisition

The untargeted metabolomics analysis was completed using an ultra-high performance liquid chromatography system (Thermo Dionex Ultimate 3000 UHPLC) coupled to an ultra-high resolution quadrupole time of flight (qTOF) mass spectrometer (Bruker Daltonics MaXis HD). A Phenomenex Kinetex column (C18 1.7 µm, 2.1 mm x 50 mm) was used for chromatographic separation. An injection volume of 5 µL was used for each sample and a flow-rate of 0.500 mL was used throughout the analysis. The mobile phase consisted of solvent A: 100% LC-MS grade water spiked with 0.1% formic acid and solvent B: 100% LC-MS grade acetonitrile spiked with 0.1% formic acid. The chromatographic gradient was as follows: 0.0–1.0 min, 5% B; 1.0– 9.0LJmin, 5–100% B; 9.0-11.0 min, 100% B; 11.0-11.5 min, 100-5% B; 11.5-12.5 min, 5% B. Data was collected with electrospray ionization in positive mode, and was saved as .d file folders.

#### Data processing

The raw .d data files were converted to mzXML format using Bruker Compass DataAnalysis 4.1 software. The resulting .mzXML file, the original .d file folders, and basic prep information sheet are stored in the UC San Diego MassIVE data repository under the accession number MSV000084550. For MS1 level feature detection, the open-source software MZmine version 2.51 (RRID:SCR_012040, https://mzmine.github.io/) was used. The parameters used are as follows: 1) Mass Detection (Centroid, Noise Level MS1 1E3, MS2 1E2); 2) ADAP Chromatogram Builder (Min Group size in # of scans=3, Group Intensity Threshold= 3E3, Min Highest Intensity=1E3, m/z tolerance 0.01 m/z or 10.0 ppm); 3) Chromatogram Deconvolution (Local Minimum Search>Chromatographic Threshold 0.01%, Minimum in RT range 0.50 min, <Minimum Relative Height 0.01%, Minimum Absolute Height 3E3, Min Ratio of Peak Top/Edge 2, Peak Duration Range 0.05-0.50 min; m/z Calculation Auto, m/z range for MS2 pairing 0.01 Da, and RT Range for MS2 Pairing 0.1 min); Isotopic Peaks Grouper (m/z Tolerance 0.01 m/z or 10.0 ppm, Retention Time Tolerance 0.3 min, Maximum Charge 4, Representative Ion Most Intense); Join Aligner (m/z Tolerance 0.01 m/z or 10.0 ppm, Weight for m/z 75, Retention Time Tolerance 0.3 min, Weight for RT 25); Gap-Filling Peak Finder (Intensity Tolerance 20%, m/z Tolerance 0.005 m/z or 10.0 ppm, Retention Time Tolerance 0.2 min). The resulting feature table was saved as a .csv file and .mgf file for use in GNPS and MetaboAnalyst.

#### Molecular networking and statistical analysis

Molecular networking was performed using the feature networking tool available through the Global Natural Products Social Molecular Networking (GNPS, RRID:SCR_019012, https://gnps.ucsd.edu/ProteoSAFe/static/gnps-splash.jsp) portal: https://gnps.ucsd.edu/ProteoSAFe/index.jsp?params=%7B%22workflow%22:%22FEATURE-BASED-MOLECULAR-NETWORKING%22,%22library_on_server%22:%22d.speclibs;%22%7D.

The annotations obtained using this workflow fall under MSI level 2 or 3 and were used for feature analysis^48^. Briefly, level 2 compounds are putatively annotated, meaning they are not identified using chemical reference standards but rather based on physical properties and/or spectral similarities to available spectral libraries (publicly available and purchased NIST17 CID). Level 3 compounds are putatively characterized classes of compounds identified similarly to level 2 compounds. The feature-based molecular networking workflow on GNPS^129^ was utilized in order to analyze the spectra associated with the feature tables produced using the open source software Mzmine version 2.51^101^ (RRID:SCR_012040, https://mzmine.github.io/). The .mgf and .csv outputs from MZmine v2.51 were used to run the workflow. The GNPS workflow parameters used were as follows: Precursor Ion Mass 0.02 Da, Fragment Ion Mass Tolerance 0.02 Da, Min Pairs Cos 0.7, Minimum Matched Fragments 6, Maximum Shift Between Precursors 500 Da, Network TopK 10, Maximum Connected Component Size (Beta) 100, and the files were row sum normalized. Default parameters were used for the rest of the settings. Visualizations and statistical analyses were performed using QIIME 2 v.2019.10^95^ (RRID:SCR_021258), MetaboAnalyst^102^ (RRID:SCR_015539, https://www.metaboanalyst.ca/) and Cytoscape v3.7.2^103^ (RRID:SCR_003032, https://cytoscape.org/). The metabolomics data from this study are available from (https://github.com/mazmanianlab/Griffiths_Yoo_et_al/tree/main/metabolomics).

### Proteome Preparation

Procedures are described in dx.doi.org/10.17504/protocols.io.bzqcp5sw.

#### Protein extraction

Mice were sacrificed 1 hour after the final C21 administration and cecal contents were isolated and resuspended in 400 μl of phosphate buffered solution and centrifuged at 20,000 xg to spin down cells and lysate. Protein was isolated from the resulting supernatant using Wessel-Flügge’s methanol/chloroform extraction method (Wessel and Flügge, 1984). Briefly, MeOH and chloroform were added to the samples at a 4:1 and 1:1 ratio, respectively. Next, dH_2_O was added at a 3:1 ratio, samples were vortexed and centrifuged at 20,000 xg. Resulting precipitated protein was collected and washed with MeOH. Precipitated protein was centrifuged and left to air dry, and stored at −20 °C until protein digestion.

#### In-solution protein digestion and desalting

Precipitated protein samples were denatured in 40 µL of 8M Urea (100 mM Tris-HCl pH 8.5). To reduce disulfide bonds, 1.25 µL of 100 mM Tris(2-carboxyethyl)Phosphine was added and incubated at room temperature (RT) for 20 minutes. Then 1.8 µL of 250 mM iodoacetamide was added and incubated at RT in the dark to alkylate cysteines. The first step of digestion was initiated by adding 1 µL of 0.1 µg/µL of lysyl endopeptidase. After 4 hours of incubation, the urea concentration was adjusted to 2 M by adding 120 µL of 100 mM Tris-HCl pH 8.5. The second step of digestion was done by adding 2.5 µL of 2µg/µL trypsin plus 1.6 µL of 100 mM CaCl_2_ and incubating overnight in the dark. Formic acid was added to stop trypsin digestion. Digested peptides were desalted by HPLC using a C8 peptide microtrap (Microm Bioresources), lyophilized, and diluted to 200 ng/µl in 0.2% formic acid prior to LC-MS/MS analysis.

#### LC-MS/MS

Samples were analyzed on a Q Exactive HF Orbitrap mass spectrometer coupled to an EASY nLC 1200 liquid chromatographic system (Thermo Scientific, San Jose, CA). Approximately 200 ng of peptides were loaded on a 50 µm I.D. × 25 cm column with a 10 µm electrospray tip (PicoFrit from New Objective, Woburn, MA) in-house-packed with ReproSil-Pur C18-AQ 1.9 µm (Dr. Maisch, Ammerbuch, Germany). Solvent A consisted of 2% MeCN in 0.2% FA and solvent B consisted of 80% MeCN in 0.2% FA. A non-linear 60 minute gradient from 2% B to 40% B was used to separate the peptides for analysis. The mass spectrometer was operated in a data-dependent mode, with MS1 scans collected from 400-1650 m/z at 60,000 resolution and MS/MS scans collected from 200-2000 m/z at 30,000 resolution. Dynamic exclusion of 45 s was used. The top 12 most abundant peptides with charge states between 2 and 5 were selected for fragmentation with normalized collision energy of 28.

#### Peptide and protein identification

Thermo .raw files were converted to .ms1 and .ms2 files using RawConverter 1.1.0.18 operating in data dependent mode and selecting for monoisotopic m/z. Tandem mass spectra (.ms2 files) were identified by a database search method using the Integrated Proteomics Pipeline 6.5.4 (IP2, Integrated Proteomics Applications, Inc., http://www.integratedproteomics.com). Briefly, databases containing forward and reverse (decoy)^130,131^ peptide sequences were generated from *in silico* trypsin digestion of either the mouse proteome (UniProt; Oct. 2, 2019)^104^ or protein sequences derived from large comprehensive public repositories (ComPIL 2.0)^105^. Tandem mass spectra were matched to peptide sequences using the ProLuCID/SEQUEST (1.4)^106,107^ software package. The validity of spectrum-peptide matches was assessed using the SEQUEST-defined parameters XCorr (cross-correlation score) and DeltaCN (normalized difference in cross-correlation scores) in the DTASelect2 (2.1.4)^108,109^ software package. Search settings were configured as follows: (1) 5 ppm precursor ion mass tolerance, (2) 10 ppm fragment ion mass tolerance, (3) 1% peptide false discovery rate, (4) 2 peptide per protein minimum, (5) 600-6000 Da precursor mass window, (6) 2 differential modifications per peptide maximum (methionine oxidation: M+15.994915 Da), (7) unlimited static modifications per peptide (cysteine carbamidomethylation: C+57.02146 Da), and (8) the search space included half- and fully-tryptic (cleavage C-terminal to K and R residues) peptide candidates with unlimited (mouse database, custom metagenomic shotgun database) or 2 missed cleavage events (ComPIL 2.0).

### Differential analysis of detected proteins using peptide-spectrum matches (spectral counts)

Detected proteins were grouped by sequence similarity into “clusters” using CD-HIT 4.8.1 (RRID:SCR_007105, http://weizhong-lab.ucsd.edu/cd-hit/)^110,111,132^ at the following similarity cut-offs: 65%, 75%, 85%, and 95%. The following is an example command line input: “cd-hit-i fastafile.fasta-o outputfile -c 0.65 -g 1 -d 0”. Tandem mass spectra identified as peptides (peptide spectrum matches, PSMs) were mapped to CD-HIT generated clusters. PSMs mapping to >1 cluster were discarded. Cluster-PSM tables were generated and differential analysis was performed in DESeq2 (1.25.13, RRID:SCR_015687, https://bioconductor.org/packages/release/bioc/html/DESeq2.html)^112^. Briefly, count data (PSMs) were modeled using the negative binomial distribution, and the mean-variance relationship was estimated. Variance was estimated using an information sharing approach whereby a single feature’s (or cluster’s) variance was estimated by taking into account variances of other clusters measured in the same experiment. Feature significance calling and ranking were performed using estimated effect sizes. Multiple testing correction was performed by the Benjamini-Hochberg method within the DESeq2 package. Volcano plots were generated in Prism (GraphPad).

### Differential analysis of detected proteins using ion intensity (precursor intensity)

Detected proteins were grouped into “clusters” by sequence similarity using CD-HIT 4.8.1^110,111,132^ at the following similarity cut-offs: 65%, 75%, 85%, and 95%. The following is an example command line input: “cd-hit-i fastafile.fasta -o outputfile -c 0.65 -g 1 -d 0”. Using the Census software package^133^ (Integrated Proteomics Pipeline 6.5.4), peptide ion intensities were calculated from .ms1 files. Peptide ion intensities were assigned to their parent peptide, then parent peptides were mapped to their appropriate CD-HIT generated clusters. Ion intensities belonging to parent peptides that mapped to >1 CD-HIT cluster were discarded. Cluster-ion intensity tables were generated.

Ion intensity data were analyzed using the Differential Enrichment analysis of Proteomics data DEP package (RRID:SCR_023090, https://bioconductor.org/packages/release/bioc/html/DEP.html)^113^ operating in R. Intensity values were automatically Log2 transformed in DEP. The cluster list was subsequently filtered with the ‘filter_proteins’ function such that clusters with missing values above a 65% threshold were discarded. Remaining intensities were further transformed by the ‘normalize_vsn’ function^134^. Missing data in remaining clusters were imputed using a mixed approach. Clusters where either the control or treatment group contained only null entries were classified as ‘missing not at random’ (MNAR) and imputed with 0 values. All other groups were treated as ‘missing at random’ (MAR) and imputed using the maximum likelihood method (’MLE’)^135^. Note that for a given cluster, missing values for treatment groups were imputed separately by treatment group. Differential expression analyses were performed on filled-in cluster-ion intensity tables using the ‘test_diff’ function^136^ and multiple testing correction was performed using the ‘add_rejections’ function.

#### Network analysis using the STRING database

Upregulated proteins with a nominal p-value < 0.2 were searched against protein-protein interactions in the STRING database^114^ (RRID:SCR_005223, http://www.string-db.org) where high confidence interactions are selected for. Briefly, the STRING database sources protein-protein interactions from primary databases consisting of genomic context predictions, high-throughput lab experiments, (conserved) co-expression, automated text mining, and previous knowledge in databases^114^.

#### Metaproteome analysis using Unipept

Upregulated tryptic, microbial peptide sequences, with fold change and nominal p-value cutoffs of >2 and <0.2, respectively, were input into Unipept (http://unipept.ugent.be)^137,138^, equating leucine and isoleucine and filtering duplicate peptides. Briefly, Unipept indexes tryptic peptide sequences from the UniProtKB and details peptides with NCBI’s taxonomic database. Lowest common ancestor was calculated for each tryptic peptide. The proteomics data from this study are available from (https://github.com/mazmanianlab/Griffiths_Yoo_et_al/tree/main/proteomics).

### 3’ mRNA sequencing

Procedures are described in dx.doi.org/10.17504/protocols.io.bzqbp5sn.

#### Tissue collection and RNA extraction

Mice were cervically dislocated and the GI tract was removed. 1 cm of tissue above and below the cecum were dissected and cleaned to represent tissue from the distal SI and proximal colon, respectively. Tissue was homogenized in TRIzol (ThermoFisher Scientific, Waltham. MA-Cat. No. 15596018) solution using a bead-based homogenizing method, and total RNA was extracted using chloroform per manufacturer’s instructions.

#### Library preparation, sequencing, and analysis

The cDNA libraries were prepared using the QuantSeq 3’mRNA-Seq Library Prep Kit FWD for Illumina (Lexogen, Greenland, NH) supplemented with UMI (unique molecular index) as per the manufacturer’s instructions. Briefly, total RNA was reverse transcribed using oligo (dT) primers. The second cDNA strand was synthesized by random priming, in a manner such that DNA polymerase is efficiently stopped when reaching the next hybridized random primer, so only the fragment closest to the 3’ end gets captured for later indexed adapter ligation and PCR amplification. UMIs were incorporated to the first 6 bases of each read, followed by 4 bases of spacer sequences. UMIs are used to eliminate possible PCR duplicates in sequencing datasets and therefore facilitate unbiased gene expression profiling. The basic principle behind the UMI deduplication step is to collapse reads with identical mapping coordinates and UMI sequences. This step helps increase the accuracy of sequencing read counts for downstream analysis of gene expression levels. The processed libraries were assessed for size distribution and concentration using the Bioanalyzer High Sensitivity DNA Kit (Agilent Technologies, Santa Clara, CA-Cat. No. 5067-4626 and −4627). Pooled libraries were diluted to 2 nM in EB buffer (Qiagen, Hilden, Germany, Cat. No. 19086) and then denatured using the Illumina protocol. The libraries were pooled and diluted to 2 nM using 10 mM Tris-HCl, pH 8.5 and then denatured using the Illumina protocol. The denatured libraries were diluted to 10 pM with pre-chilled hybridization buffer and loaded onto an Illumina MiSeq v3 flow cell for 150 cycles using a single-read recipe according to the manufacturer’s instructions. Single-end 75 bp reads (max 4.5M reads) were obtained. De-multiplexed sequencing reads were generated using Illumina BaseSpace.

UMI specific workflows that were developed and distributed by Lexogen were used to extract reads that are free from PCR artifacts (i.e., deduplication). First, the umi2index tool was used to add the 6 nucleotide UMI sequence to the identifier of each read and trim the UMI from the start of each read. This generated a new FASTQ file, which was then processed through trimming and alignment. Second, after the quality and polyA trimming by BBDuk^115^ (Bestus Bioinformaticus Duk, RRID:SCR_016969, https://jgi.doe.gov/data-and-tools/bbtools/bb-tools-user-guide/bbduk-guide/) and alignment by HISAT2 (version 2.1.0, RRID:SCR_015530, https://daehwankimlab.github.io/hisat2/)^116^, the mapped reads were collapsed according to the UMI sequence of each read. Reads were collapsed if they had the same mapping coordinates (CIGAR string) and identical UMI sequences. Collapsing reads in this manner removes PCR duplicates. Read counts were calculated using HTSeq (RRID:SCR_005514, https://htseq.readthedocs.io/en/release_0.9.1/)^117^ by supplementing Ensembl gene annotation (GRCm38.78). Raw read counts were run through ShinySeq to obtain differentially expressed genes and downstream gene ontology analyses^139^. The RNAseq data from this study are available from (https://github.com/mazmanianlab/Griffiths_Yoo_et_al/tree/main/RNAseq).

### Whole Gut Transit Time, Fecal Water Content, and Fecal Output

Procedures are described in dx.doi.org/<u>10.17504/protocols.io.36wgq3p1xlk5/v1</u>. 6% (w/v) carmine red (Sigma-Aldrich, St. Louis, MO) with 0.5% methylcellulose (Sigma-Aldrich) was dissolved and autoclaved prior to use. ChAT-Cre and TH-Cre mice were administered C21 (3 mg/kg) intraperitoneally, and subsequently orally gavaged with 150 µL of carmine red solution. Mice were single-housed with no bedding for the duration of the experiment, and animals were not fasted beforehand. Over the 5 hours following gavage, the time of expulsion was recorded for each fecal pellet, and each pellet was collected in pre-weighed, 1.5 mL microcentrifuge tube. Each pellet collected was checked for the presence of carmine red, and the time of initial carmine red pellet expulsion was recorded as GI transit time. The mass of collected fecal pellets was determined, and pellets were left to dry in an 80 °C oven for 2 days before weighing the desiccated pellets and calculating the pellets’ initial water content. Fecal output rate for each mouse was calculated as the total number of pellets expelled during the 5 hour time course post-C21 administration divided by the time the last fecal pellet was expelled.

### Colonic Migrating Motor Complexes in *Ex Vivo* Intestinal Preparations

Procedures are described in dx.doi.org/<u>10.17504/protocols.io.n92ldm61nl5b/v1</u>. Intact colons were dissected from cervically-dislocated ChAT-Cre and TH-Cre mice, flushed and placed in pre-oxygenated (95% O_2_, 5% CO_2_) Krebs-Henseleit solution at RT. Proximal and distal ends were cannulated to 2 mm diameter tubes and secured in the center of an organ bath with continuously oxygenated Krebs-Henseleit solution at 37 °C. Syringe pumps were connected to the inlet and outlet tubes to maintain a flow of solution at a rate of 500 µL/min through the colon. The system was allowed to equilibrate for 30 minutes before recording. Baseline recordings were taken for 30 minutes, then the Krebs solution in the organ bath was briefly removed, mixed with C21 to a final concentration of 2 µM, and replaced in the organ bath. Recordings were taken for another 30 minutes.

### Quantification and Statistical Analysis

Statistical methodologies and software used for performing various types of multi-omic analysis in this work are cited where appropriate in the STAR Methods text. The p-value calculations for viral transduction, microbiome differences, gastrointestinal function, and animal welfare were done using Prism GraphPad v.9.2.0. The specific statistical test used for each figure is denoted in the figure legends. Error bars represent the standard error of the mean unless otherwise stated.

## Supplemental Video and Table Legends

**Table S1 - Extended GNPS Annotations of Metabolite Network Nodes (Related to Figure 3).**

Extended annotations for networked (bold) MS/MS spectra from luminal cecal contents of activated TH^+^ and ChAT^+^ mice in Figures 3C and 3D, respectively.

Source Data Table S1 https://github.com/mazmanianlab/Griffiths_Yoo_et_al/blob/main/metabolomics/Table_S1-Extended_GNPS_annotations_of_Metabolite_Network_Notes_related_to_figure_3.xlsx

**Table S2 - Gene Set Enrichment Analysis of Gene Ontology (GO) Terms (Related to Figure 5).**

Gene set enrichment analysis of gene ontology (GO) terms that are upregulated in the distal SI and proximal colon of activated ChAT^+^ and TH^+^ mice.

Source Data Table S2 https://github.com/mazmanianlab/Griffiths_Yoo_et_al/blob/main/RNAseq/Table_S2-Gene_Set_Enrichment_Analysis_of_GO_related_to_figure_5.xlsx

**Table S3 – Genes Related to T helper responses (Related to Figure 5).**

List of genes in the transcriptomics dataset associated with T helper responses. Source Data Table S3 https://github.com/mazmanianlab/Griffiths_Yoo_et_al/blob/main/RNAseq/Table_S3-Genes_Related_to_T_Helper_Responses.xlsx

**Table S4 – Annotations of Colonic Migrating Motor Complexes (Related to Figure 6)** Video annotations of CMMCs in *ex vivo* colonic preparations from ChAT^+^ and TH^+^ mice at baseline and during activation.

Source Data Table S4 https://github.com/mazmanianlab/Griffiths_Yoo_et_al/blob/main/ex_vivo_motility/Table_S4%E2%80%93Annotations_of_Colonic_Migrating_Motor_Complexes_Related_to_figure_6_.xlsx

**Video S1 – Calcium Imaging of ChAT^+^ activated Gut Neurons (Related to Figure S5)** Video of *in vivo* calcium imaging of GCaMP6f-expressing ChAT^+^ activated neurons in the myenteric plexus at 5 Hz following C21 administration.

Source Data Video S1 https://github.com/mazmanianlab/Griffiths_Yoo_et_al/blob/main/ENS%20quantification/Video%20Supplement%201-%20GCaMP6F%20at%205Hz.avi

## REFERENCES

1. Furness JB. The Enteric Nervous System. Wiley; 2008. https://books.google.com/books?id=pvkpdNHhI6cC

2. Rao M, Gershon MD. The bowel and beyond: the enteric nervous system in neurological disorders. Nat Rev Gastroenterol Hepatol. 2016;13(9):517–528. doi:10.1038/nrgastro.2016.107

3. Grundy D, Brookes S. Neural Control of Gastrointestinal Function. Morgan & Claypool Life Science Publishers; 2011. https://books.google.com/books?id=NV2BMJk6XHwC

4. Furness JB, Callaghan BP, Rivera LR, Cho HJ. The enteric nervous system and gastrointestinal innervation: integrated local and central control. Adv Exp Med Biol. 2014;817:39–71. doi:10.1007/978-1-4939-0897-4_3

5. Schneider S, Wright CM, Heuckeroth RO. Unexpected Roles for the Second Brain: Enteric Nervous System as Master Regulator of Bowel Function. Annu Rev Physiol. 2019;81:235–259. doi:10.1146/annurev-physiol-021317-121515

6. Gabanyi I, Muller PA, Feighery L, Oliveira TY, Costa-Pinto FA, Mucida D. Neuro-immune Interactions Drive Tissue Programming in Intestinal Macrophages. Cell. 2016;164(3):378–391. doi:10.1016/j.cell.2015.12.023

7. Muller PA, Koscsó B, Rajani GM, et al. Crosstalk between muscularis macrophages and enteric neurons regulates gastrointestinal motility. Cell. 2014;158(2):300–313. doi:10.1016/j.cell.2014.04.050

8. Bravo JA, Forsythe P, Chew MV, et al. Ingestion of Lactobacillus strain regulates emotional behavior and central GABA receptor expression in a mouse via the vagus nerve. Proc Natl Acad Sci. 2011;108(38):16050–16055. doi:10.1073/pnas.1102999108

9. Kaelberer MM, Buchanan KL, Klein ME, et al. A gut-brain neural circuit for nutrient sensory transduction. Science. 2018;361(6408). doi:10.1126/science.aat5236

10. Jarret A, Jackson R, Duizer C, et al. Enteric Nervous System-Derived IL-18 Orchestrates Mucosal Barrier Immunity. Cell. 2020;180(1):50–63.e12. doi:10.1016/j.cell.2019.12.016

11. Seillet C, Luong K, Tellier J, et al. The neuropeptide VIP confers anticipatory mucosal immunity by regulating ILC3 activity. Nat Immunol. 2020;21(2):168–177. doi:10.1038/s41590-019-0567-y

12. Talbot J, Hahn P, Kroehling L, Nguyen H, Li D, Littman DR. Feeding-dependent VIP neuron–ILC3 circuit regulates the intestinal barrier. Nature. 2020;579(7800):575-580. doi:10.1038/s41586-020-2039-9

13. Lai NY, Musser MA, Pinho-Ribeiro FA, et al. Gut-Innervating Nociceptor Neurons Regulate Peyer’s Patch Microfold Cells and SFB Levels to Mediate Salmonella Host Defense. Cell. 2020;180(1):33–49.e22. doi:10.1016/j.cell.2019.11.014

14. Matheis F, Muller PA, Graves CL, et al. Adrenergic Signaling in Muscularis Macrophages Limits Infection-Induced Neuronal Loss. Cell. 2020;180(1):64–78.e16. doi:10.1016/j.cell.2019.12.002

15. Nezami BG, Srinivasan S. Enteric nervous system in the small intestine: pathophysiology and clinical implications. Curr Gastroenterol Rep. 2010;12(5):358–365. doi:10.1007/s11894-010-0129-9

16. Qu ZD, Thacker M, Castelucci P, Bagyánszki M, Epstein ML, Furness JB. Immunohistochemical analysis of neuron types in the mouse small intestine. Cell Tissue Res. 2008;334(2):147–161. doi:10.1007/s00441-008-0684-7

17. Furness JB. The enteric nervous system and neurogastroenterology. Nat Rev Gastroenterol Hepatol. 2012;9(5):286–294. doi:10.1038/nrgastro.2012.32

18. Hennig GW, Gould TW, Koh SD, et al. Use of Genetically Encoded Calcium Indicators (GECIs) Combined with Advanced Motion Tracking Techniques to Examine the Behavior of Neurons and Glia in the Enteric Nervous System of the Intact Murine Colon. Front Cell Neurosci. 2015;9:436. doi:10.3389/fncel.2015.00436

19. Niesler B, Kuerten S, Demir IE, Schäfer KH. Disorders of the enteric nervous system — a holistic view. Nat Rev Gastroenterol Hepatol. 2021;18(6):393–410. doi:10.1038/s41575-020-00385-2

20. Nezami BG, Srinivasan S. Enteric nervous system in the small intestine: pathophysiology and clinical implications. Curr Gastroenterol Rep. 2010;12(5):358–365. doi:10.1007/s11894-010-0129-9

21. Qu ZD, Thacker M, Castelucci P, Bagyánszki M, Epstein ML, Furness JB. Immunohistochemical analysis of neuron types in the mouse small intestine. Cell Tissue Res. 2008;334(2):147–161. doi:10.1007/s00441-008-0684-7

22. Lott EL, Jones EB. Cholinergic Toxicity. In: StatPearls. StatPearls Publishing; 2023.

23. Monane M, Avorn J, Beers MH, Everitt DE. Anticholinergic drug use and bowel function in nursing home patients. Arch Intern Med. 1993;153(5):633–638.

24. Fung C, Koussoulas K, Unterweger P, Allen AM, Bornstein JC, Foong JPP. Cholinergic Submucosal Neurons Display Increased Excitability Following in Vivo Cholera Toxin Exposure in Mouse Ileum. Front Physiol. 2018;9:260. doi:10.3389/fphys.2018.00260

25. Wang H, Foong JPP, Harris NL, Bornstein JC. Enteric neuroimmune interactions coordinate intestinal responses in health and disease. Mucosal Immunol. 2022;15(1):27–39. doi:10.1038/s41385-021-00443-1

26. Li ZS, Schmauss C, Cuenca A, Ratcliffe E, Gershon MD. Physiological modulation of intestinal motility by enteric dopaminergic neurons and the D2 receptor: analysis of dopamine receptor expression, location, development, and function in wild-type and knock-out mice. J Neurosci Off J Soc Neurosci. 2006;26(10):2798–2807. doi:10.1523/JNEUROSCI.4720-05.2006

27. Baumuratov AS, Antony PMA, Ostaszewski M, et al. Enteric neurons from Parkinson’s disease patients display ex vivo aberrations in mitochondrial structure. Sci Rep. 2016;6(1):33117. doi:10.1038/srep33117

28. McQuade RM, Singleton LM, Wu H, et al. The association of enteric neuropathy with gut phenotypes in acute and progressive models of Parkinson’s disease. Sci Rep. 2021;11(1):7934. doi:10.1038/s41598-021-86917-5

29. Mittal R, Debs LH, Patel AP, et al. Neurotransmitters: The Critical Modulators Regulating Gut-Brain Axis. J Cell Physiol. 2017;232(9):2359–2372. doi:10.1002/jcp.25518

30. Chan KY, Jang MJ, Yoo BB, et al. Engineered AAVs for efficient noninvasive gene delivery to the central and peripheral nervous systems. Nat Neurosci. 2017;20(8):1172–1179. doi:10.1038/nn.4593

31. Wess J, Nakajima K, Jain S. Novel designer receptors to probe GPCR signaling and physiology. Trends Pharmacol Sci. 2013;34(7):385–392. doi:10.1016/j.tips.2013.04.006

32. Furness JB. The organisation of the autonomic nervous system: peripheral connections. Auton Neurosci Basic Clin. 2006;130(1-2):1–5. doi:10.1016/j.autneu.2006.05.003

33. Hamnett R, Dershowitz LB, Sampathkumar V, et al. Regional cytoarchitecture of the adult and developing mouse enteric nervous system. Curr Biol CB. 2022;32(20):4483–4492.e5. doi:10.1016/j.cub.2022.08.030

34. Hama H, Hioki H, Namiki K, et al. ScaleS: an optical clearing palette for biological imaging. Nat Neurosci. 2015;18(10):1518–1529. doi:10.1038/nn.4107

35. Treweek JB, Chan KY, Flytzanis NC, et al. Whole-body tissue stabilization and selective extractions via tissue-hydrogel hybrids for high-resolution intact circuit mapping and phenotyping. Nat Protoc. 2015;10(11):1860–1896. doi:10.1038/nprot.2015.122

36. Yang B, Treweek JB, Kulkarni RP, et al. Single-cell phenotyping within transparent intact tissue through whole-body clearing. Cell. 2014;158(4):945–958. doi:10.1016/j.cell.2014.07.017

37. Deverman BE, Pravdo PL, Simpson BP, et al. Cre-dependent selection yields AAV variants for widespread gene transfer to the adult brain. Nat Biotechnol. 2016;34(2):204–209. doi:10.1038/nbt.3440

38. Chan KY, Jang MJ, Yoo BB, et al. Engineered AAVs for efficient noninvasive gene delivery to the central and peripheral nervous systems. Nat Neurosci. 2017;20(8):1172–1179. doi:10.1038/nn.4593

39. Haenraets K, Foster E, Johannssen H, et al. Spinal nociceptive circuit analysis with recombinant adeno-associated viruses: the impact of serotypes and promoters. J Neurochem. 2017;142(5):721–733. doi:10.1111/jnc.14124

40. Jakob MO, Kofoed-Branzk M, Deshpande D, Murugan S, Klose CSN. An Integrated View on Neuronal Subsets in the Peripheral Nervous System and Their Role in Immunoregulation. Front Immunol. 2021;12:679055. doi:10.3389/fimmu.2021.679055

41. Tavares-Ferreira D, Shiers S, Ray PR, et al. Spatial transcriptomics of dorsal root ganglia identifies molecular signatures of human nociceptors. Sci Transl Med. 2022;14(632):eabj8186. doi:10.1126/scitranslmed.abj8186

42. Challis RC, Kumar SR, Chan KY, et al. Systemic AAV vectors for widespread and targeted gene delivery in rodents. Nat Protoc. 2019;14(2):379–414. doi:10.1038/s41596-018-0097-3

43. Kaestner CL, Smith EH, Peirce SG, Hoover DB. Immunohistochemical analysis of the mouse celiac ganglion: An integrative relay station of the peripheral nervous system. J Comp Neurol. 2019;527(16):2742–2760. doi:10.1002/cne.24705

44. Browning KN, Travagli RA. Central nervous system control of gastrointestinal motility and secretion and modulation of gastrointestinal functions. Compr Physiol. 2014;4(4):1339–1368. doi:10.1002/cphy.c130055

45. Thompson KJ, Khajehali E, Bradley SJ, et al. DREADD Agonist 21 Is an Effective Agonist for Muscarinic-Based DREADDs in Vitro and in Vivo. ACS Pharmacol Transl Sci. 2018;1(1):61–72. doi:10.1021/acsptsci.8b00012

46. Segata N, Izard J, Waldron L, et al. Metagenomic biomarker discovery and explanation. Genome Biol. 2011;12(6):R60. doi:10.1186/gb-2011-12-6-r60

47. Wang M, Carver JJ, Phelan VV, et al. Sharing and community curation of mass spectrometry data with Global Natural Products Social Molecular Networking. Nat Biotechnol. 2016;34(8):828–837. doi:10.1038/nbt.3597

48. Sumner LW, Amberg A, Barrett D, et al. Proposed minimum reporting standards for chemical analysis Chemical Analysis Working Group (CAWG) Metabolomics Standards Initiative (MSI). Metabolomics Off J Metabolomic Soc. 2007;3(3):211–221. doi:10.1007/s11306-007-0082-2

49. Aries V, Crowther JS, Drasar BS, Hill MJ. Degradation of bile salts by human intestinal bacteria. Gut. 1969;10(7):575–576. doi:10.1136/gut.10.7.575

50. Sakai K, Makino T, Kawai Y, Mutai M. Intestinal microflora and bile acids. Effect of bile acids on the distribution of microflora and bile acid in the digestive tract of the rat. Microbiol Immunol. 1980;24(3):187–196. doi:10.1111/j.1348-0421.1980.tb00578.x

51. Albenberg LG, Wu GD. Diet and the intestinal microbiome: associations, functions, and implications for health and disease. Gastroenterology. 2014;146(6):1564–1572. doi:10.1053/j.gastro.2014.01.058

52. Jia L, Betters JL, Yu L. Niemann-pick C1-like 1 (NPC1L1) protein in intestinal and hepatic cholesterol transport. Annu Rev Physiol. 2011;73:239–259. doi:10.1146/annurev-physiol-012110-142233

53. Rodríguez-Piñeiro AM, Bergström JH, Ermund A, et al. Studies of mucus in mouse stomach, small intestine, and colon. II. Gastrointestinal mucus proteome reveals Muc2 and Muc5ac accompanied by a set of core proteins. Am J Physiol Gastrointest Liver Physiol. 2013;305(5):G348-356. doi:10.1152/ajpgi.00047.2013

54. Matsushima S, Hori S, Matsuda M. Conversion of 4-aminobutyraldehyde to gamma-aminobutyric acid in striatum treated with semicarbazide and kainic acid. Neurochem Res. 1986;11(9):1313–1319. doi:10.1007/BF00966125

55. Li W, Yu G, Liu Y, Sha L. Intrapancreatic Ganglia and Neural Regulation of Pancreatic Endocrine Secretion. Front Neurosci. 2019;13. doi:10.3389/fnins.2019.00021

56. Donowitz M, Singh S, Salahuddin FF, et al. Proteome of murine jejunal brush border membrane vesicles. J Proteome Res. 2007;6(10):4068–4079. doi:10.1021/pr0701761

57. McConnell RE, Benesh AE, Mao S, Tabb DL, Tyska MJ. Proteomic analysis of the enterocyte brush border. Am J Physiol Gastrointest Liver Physiol. 2011;300(5):G914–926. doi:10.1152/ajpgi.00005.2011

58. Latgé JP. The cell wall: a carbohydrate armour for the fungal cell. Mol Microbiol. 2007;66(2):279–290. doi:10.1111/j.1365-2958.2007.05872.x

59. Wu YE, Pan L, Zuo Y, Li X, Hong W. Detecting Activated Cell Populations Using Single-Cell RNA-Seq. Neuron. 2017;96(2):313–329.e6. doi:10.1016/j.neuron.2017.09.026

60. Bahrami S, Drabløs F. Gene regulation in the immediate-early response process. Adv Biol Regul. 2016;62:37–49. doi:10.1016/j.jbior.2016.05.001

61. Ramirez-Carrozzi VR, Braas D, Bhatt DM, et al. A unifying model for the selective regulation of inducible transcription by CpG islands and nucleosome remodeling. Cell. 2009;138(1):114–128. doi:10.1016/j.cell.2009.04.020

62. Miano JM, Vlasic N, Tota RR, Stemerman MB. Smooth muscle cell immediate-early gene and growth factor activation follows vascular injury. A putative in vivo mechanism for autocrine growth. Arterioscler Thromb J Vasc Biol. 1993;13(2):211–219. doi:10.1161/01.atv.13.2.211

63. Flandez M, Guilmeau S, Blache P, Augenlicht LH. KLF4 regulation in intestinal epithelial cell maturation. Exp Cell Res. 2008;314(20):3712–3723. doi:10.1016/j.yexcr.2008.10.004

64. Furness JB. The enteric nervous system and neurogastroenterology. Nat Rev Gastroenterol Hepatol. 2012;9(5):286–294. doi:10.1038/nrgastro.2012.32

65. Fung C, Koussoulas K, Unterweger P, Allen AM, Bornstein JC, Foong JPP. Cholinergic Submucosal Neurons Display Increased Excitability Following in Vivo Cholera Toxin Exposure in Mouse Ileum. Front Physiol. 2018;9:260. doi:10.3389/fphys.2018.00260

66. Monane M, Avorn J, Beers MH, Everitt DE. Anticholinergic drug use and bowel function in nursing home patients. Arch Intern Med. 1993;153(5):633–638.

67. Johnson CD, Barlow-Anacker AJ, Pierre JF, et al. Deletion of choline acetyltransferase in enteric neurons results in postnatal intestinal dysmotility and dysbiosis. FASEB J Off Publ Fed Am Soc Exp Biol. 2018;32(9):4744–4752. doi:10.1096/fj.201701474RR

68. Everard A, Belzer C, Geurts L, et al. Cross-talk between Akkermansia muciniphila and intestinal epithelium controls diet-induced obesity. Proc Natl Acad Sci. 2013;110(22):9066–9071. doi:10.1073/pnas.1219451110

69. Plovier H, Everard A, Druart C, et al. A purified membrane protein from Akkermansia muciniphila or the pasteurized bacterium improves metabolism in obese and diabetic mice. Nat Med. 2017;23(1):107–113. doi:10.1038/nm.4236

70. Cekanaviciute E, Yoo BB, Runia TF, et al. Gut bacteria from multiple sclerosis patients modulate human T cells and exacerbate symptoms in mouse models. Proc Natl Acad Sci U S A. 2017;114(40):10713–10718. doi:10.1073/pnas.1711235114

71. Jangi S, Gandhi R, Cox LM, et al. Alterations of the human gut microbiome in multiple sclerosis. Nat Commun. 2016;7(1):12015. doi:10.1038/ncomms12015

72. Olson CA, Vuong HE, Yano JM, Liang QY, Nusbaum DJ, Hsiao EY. The Gut Microbiota Mediates the Anti-Seizure Effects of the Ketogenic Diet. Cell. 2018;173(7):1728–1741.e13. doi:10.1016/j.cell.2018.04.027

73. de Aguiar Vallim TQ, Tarling EJ, Edwards PA. Pleiotropic roles of bile acids in metabolism. Cell Metab. 2013;17(5):657–669. doi:10.1016/j.cmet.2013.03.013

74. Kirwan WO, Smith AN, Mitchell WD, Falconer JD, Eastwood MA. Bile acids and colonic motility in the rabbit and the human. Gut. 1975;16(11):894–902. doi:10.1136/gut.16.11.894

75. Watanabe M, Houten SM, Mataki C, et al. Bile acids induce energy expenditure by promoting intracellular thyroid hormone activation. Nature. 2006;439(7075):484-489. doi:10.1038/nature04330

76. Fiorucci S, Biagioli M, Zampella A, Distrutti E. Bile Acids Activated Receptors Regulate Innate Immunity. Front Immunol. 2018;9:1853. doi:10.3389/fimmu.2018.01853

77. McMillin M, DeMorrow S. Effects of bile acids on neurological function and disease. FASEB J Off Publ Fed Am Soc Exp Biol. 2016;30(11):3658–3668. doi:10.1096/fj.201600275R

78. Begley M, Sleator RD, Gahan CGM, Hill C. Contribution of three bile-associated loci, bsh, pva, and btlB, to gastrointestinal persistence and bile tolerance of Listeria monocytogenes. Infect Immun. 2005;73(2):894–904. doi:10.1128/IAI.73.2.894-904.2005

79. Delpino MV, Marchesini MI, Estein SM, et al. A bile salt hydrolase of Brucella abortus contributes to the establishment of a successful infection through the oral route in mice. Infect Immun. 2007;75(1):299–305. doi:10.1128/IAI.00952-06

80. Hofmann AF, Eckmann L. How bile acids confer gut mucosal protection against bacteria. Proc Natl Acad Sci U S A. 2006;103(12):4333–4334. doi:10.1073/pnas.0600780103

81. Jones BV, Begley M, Hill C, Gahan CGM, Marchesi JR. Functional and comparative metagenomic analysis of bile salt hydrolase activity in the human gut microbiome. Proc Natl Acad Sci. 2008;105(36):13580–13585. doi:10.1073/pnas.0804437105

82. Sannasiddappa TH, Lund PA, Clarke SR. In Vitro Antibacterial Activity of Unconjugated and Conjugated Bile Salts on Staphylococcus aureus. Front Microbiol. 2017;8:1581. doi:10.3389/fmicb.2017.01581

83. Drokhlyansky E, Smillie CS, Van Wittenberghe N, et al. The Human and Mouse Enteric Nervous System at Single-Cell Resolution. Cell. 2020;182(6):1606–1622.e23. doi:10.1016/j.cell.2020.08.003

84. Bhavsar AS, Verma S, Lamba R, Lall CG, Koenigsknecht V, Rajesh A. Abdominal manifestations of neurologic disorders. Radiogr Rev Publ Radiol Soc N Am Inc. 2013;33(1):135–153. doi:10.1148/rg.331125097

85. Cersosimo MG, Raina GB, Pecci C, et al. Gastrointestinal manifestations in Parkinson’s disease: prevalence and occurrence before motor symptoms. J Neurol. 2013;260(5):1332–1338. doi:10.1007/s00415-012-6801-2

86. Del Giudice E, Staiano A, Capano G, et al. Gastrointestinal manifestations in children with cerebral palsy. Brain Dev. 1999;21(5):307–311. doi:10.1016/s0387-7604(99)00025-x

87. Pfeiffer RF. Gastrointestinal dysfunction in Parkinson’s disease. Lancet Neurol. 2003;2(2):107–116. doi:10.1016/s1474-4422(03)00307-7

88. Valicenti-McDermott MD, McVicar K, Cohen HJ, Wershil BK, Shinnar S. Gastrointestinal symptoms in children with an autism spectrum disorder and language regression. Pediatr Neurol. 2008;39(6):392–398. doi:10.1016/j.pediatrneurol.2008.07.019

89. Rajendran PS, Challis RC, Fowlkes CC, et al. Identification of peripheral neural circuits that regulate heart rate using optogenetic and viral vector strategies. Nat Commun. 2019;10(1):1944. doi:10.1038/s41467-019-09770-1

90. Roy A, Guatimosim S, Prado VF, Gros R, Prado MAM. Cholinergic activity as a new target in diseases of the heart. Mol Med Camb Mass. 2015;20(1):527–537. doi:10.2119/molmed.2014.00125

91. Mohanta SK, Yin C, Weber C, et al. Cardiovascular Brain Circuits. Circ Res. 2023;132(11):1546-1565. doi:10.1161/CIRCRESAHA.123.322791

92. Finneran DJ, Njoku IP, Flores-Pazarin D, et al. Toward Development of Neuron Specific Transduction After Systemic Delivery of Viral Vectors. Front Neurol. 2021;12. doi:10.3389/fneur.2021.685802

93. Kügler S, Kilic E, Bähr M. Human synapsin 1 gene promoter confers highly neuron-specific long-term transgene expression from an adenoviral vector in the adult rat brain depending on the transduced area. Gene Ther. 2003;10(4):337–347. doi:10.1038/sj.gt.3301905

94. Moran GW, Leslie FC, Levison SE, Worthington J, McLaughlin JT. Enteroendocrine cells: neglected players in gastrointestinal disorders? Ther Adv Gastroenterol. 2008;1(1):51–60. doi:10.1177/1756283X08093943

95. Bolyen E, Rideout JR, Dillon MR, et al. Reproducible, interactive, scalable and extensible microbiome data science using QIIME 2. Nat Biotechnol. 2019;37(8):852–857. doi:10.1038/s41587-019-0209-9

96. Lindeberg J, Usoskin D, Bengtsson H, et al. Transgenic expression of Cre recombinase from the tyrosine hydroxylase locus. Genes N Y N 2000. 2004;40(2):67-73. doi:10.1002/gene.20065

97. Schindelin J, Arganda-Carreras I, Frise E, et al. Fiji: an open-source platform for biological-image analysis. Nat Methods. 2012;9(7):676–682. doi:10.1038/nmeth.2019

98. Langmead B, Salzberg SL. Fast gapped-read alignment with Bowtie 2. Nat Methods. 2012;9(4):357–359. doi:10.1038/nmeth.1923

99. Franzosa EA, McIver LJ, Rahnavard G, et al. Species-level functional profiling of metagenomes and metatranscriptomes. Nat Methods. 2018;15(11):962–968. doi:10.1038/s41592-018-0176-y

100. Suzek BE, Huang H, McGarvey P, Mazumder R, Wu CH. UniRef: comprehensive and non-redundant UniProt reference clusters. Bioinforma Oxf Engl. 2007;23(10):1282–1288. doi:10.1093/bioinformatics/btm098

101. Pluskal T, Castillo S, Villar-Briones A, Orešič M. MZmine 2: Modular framework for processing, visualizing, and analyzing mass spectrometry-based molecular profile data. BMC Bioinformatics. 2010;11(1):395. doi:10.1186/1471-2105-11-395

102. Xia J, Psychogios N, Young N, Wishart DS. MetaboAnalyst: a web server for metabolomic data analysis and interpretation. Nucleic Acids Res. 2009;37(Web Server issue):W652-660. doi:10.1093/nar/gkp356

103. Shannon P, Markiel A, Ozier O, et al. Cytoscape: a software environment for integrated models of biomolecular interaction networks. Genome Res. 2003;13(11):2498–2504. doi:10.1101/gr.1239303

104. The UniProt Consortium. UniProt: a worldwide hub of protein knowledge. Nucleic Acids Res. 2019;47(D1):D506–D515. doi:10.1093/nar/gky1049

105. Park SKR, Jung T, Thuy-Boun PS, Wang AY, Yates JRI, Wolan DW. ComPIL 2.0: An Updated Comprehensive Metaproteomics Database. J Proteome Res. 2019;18(2):616–622. doi:10.1021/acs.jproteome.8b00722

106. Xu T, Venable JD, Park SK, et al. ProLuCID, a fast and sensitive tandem mass spectra-based protein identification program. In: Molecular & Cellular Proteomics. Vol 5. AMER SOC BIOCHEMISTRY MOLECULAR BIOLOGY INC 9650 ROCKVILLE PIKE, BETHESDA…; 2006:S174-S174.

107. Xu T, Park SK, Venable JD, et al. ProLuCID: An improved SEQUEST-like algorithm with enhanced sensitivity and specificity. J Proteomics. 2015;129:16–24. doi:10.1016/j.jprot.2015.07.001

108. Cociorva D, L Tabb D, Yates JR. Validation of tandem mass spectrometry database search results using DTASelect. Curr Protoc Bioinforma. 2007;Chapter 13:Unit 13.4. doi:10.1002/0471250953.bi1304s16

109. Tabb DL, McDonald WH, Yates JR. DTASelect and Contrast:L Tools for Assembling and Comparing Protein Identifications from Shotgun Proteomics. J Proteome Res. 2002;1(1):21–26. doi:10.1021/pr015504q

110. Fu L, Niu B, Zhu Z, Wu S, Li W. CD-HIT: accelerated for clustering the next-generation sequencing data. Bioinforma Oxf Engl. 2012;28(23):3150–3152. doi:10.1093/bioinformatics/bts565

111. Li W, Jaroszewski L, Godzik A. Clustering of highly homologous sequences to reduce the size of large protein databases. Bioinforma Oxf Engl. 2001;17(3):282–283. doi:10.1093/bioinformatics/17.3.282

112. Love MI, Huber W, Anders S. Moderated estimation of fold change and dispersion for RNA-seq data with DESeq2. Genome Biol. 2014;15(12):550. doi:10.1186/s13059-014-0550-8

113. Zhang X, Smits AH, van Tilburg GB, Ovaa H, Huber W, Vermeulen M. Proteome-wide identification of ubiquitin interactions using UbIA-MS. Nat Protoc. 2018;13(3):530–550. doi:10.1038/nprot.2017.147

114. Szklarczyk D, Gable AL, Lyon D, et al. STRING v11: protein-protein association networks with increased coverage, supporting functional discovery in genome-wide experimental datasets. Nucleic Acids Res. 2019;47(D1):D607-D613. doi:10.1093/nar/gky1131

115. Bushnell B, Rood J, Singer E. BBMerge - Accurate paired shotgun read merging via overlap. PloS One. 2017;12(10):e0185056. doi:10.1371/journal.pone.0185056

116. Kim D, Langmead B, Salzberg SL. HISAT: a fast spliced aligner with low memory requirements. Nat Methods. 2015;12(4):357–360. doi:10.1038/nmeth.3317

117. Anders S, Pyl PT, Huber W. HTSeq--a Python framework to work with high-throughput sequencing data. Bioinforma Oxf Engl. 2015;31(2):166–169. doi:10.1093/bioinformatics/btu638

118. Marotz C, Amir A, Humphrey G, Gaffney J, Gogul G, Knight R. DNA extraction for streamlined metagenomics of diverse environmental samples. BioTechniques. 2017;62(6):290–293. doi:10.2144/000114559

119. Glenn TC, Nilsen RA, Kieran TJ, et al. Adapterama I: universal stubs and primers for 384 unique dual-indexed or 147,456 combinatorially-indexed Illumina libraries (iTru & iNext). PeerJ. 2019;7:e7755. doi:10.7717/peerj.7755

120. Costello M, Fleharty M, Abreu J, et al. Characterization and remediation of sample index swaps by non-redundant dual indexing on massively parallel sequencing platforms. BMC Genomics. 2018;19(1):332. doi:10.1186/s12864-018-4703-0

121. Sinha R, Stanley GM, Gulati GS, et al. Index switching causes “spreading-of-signal” among multiplexed samples in Illumina HiSeq 4000 DNA sequencing. bioRxiv. Published online 2017. https://api.semanticscholar.org/CorpusID:10764771

122. Gonzalez A, Navas-Molina JA, Kosciolek T, et al. Qiita: rapid, web-enabled microbiome meta-analysis. Nat Methods. 2018;15(10):796–798. doi:10.1038/s41592-018-0141-9

123. Didion JP, Martin M, Collins FS. Atropos: specific, sensitive, and speedy trimming of sequencing reads. PeerJ. 2017;5:e3720. doi:10.7717/peerj.3720

124. Zhu Q, Mai U, Pfeiffer W, et al. Phylogenomics of 10,575 genomes reveals evolutionary proximity between domains Bacteria and Archaea. Nat Commun. 2019;10(1):5477. doi:10.1038/s41467-019-13443-4

125. Hillmann Benjamin, Al-Ghalith Gabriel A., Shields-Cutler Robin R., et al. Evaluating the Information Content of Shallow Shotgun Metagenomics. mSystems. 2018;3(6):10.1128/msystems.00069-18. doi:10.1128/msystems.00069-18

126. Martino Cameron, Morton James T., Marotz Clarisse A., et al. A Novel Sparse Compositional Technique Reveals Microbial Perturbations. mSystems. 2019;4(1):10.1128/msystems.00016-19. doi:10.1128/msystems.00016-19

127. Truong DT, Franzosa EA, Tickle TL, et al. MetaPhlAn2 for enhanced metagenomic taxonomic profiling. Nat Methods. 2015;12(10):902–903. doi:10.1038/nmeth.3589

128. Caspi R, Billington R, Keseler IM, et al. The MetaCyc database of metabolic pathways and enzymes - a 2019 update. Nucleic Acids Res. 2020;48(D1):D445–D453. doi:10.1093/nar/gkz862

129. Nothias LF, Petras D, Schmid R, et al. Feature-based molecular networking in the GNPS analysis environment. Nat Methods. 2020;17(9):905–908. doi:10.1038/s41592-020-0933-6

130. Elias JE, Gygi SP. Target-decoy search strategy for increased confidence in large-scale protein identifications by mass spectrometry. Nat Methods. 2007;4(3):207–214. doi:10.1038/nmeth1019

131. Peng J, Elias JE, Thoreen CC, Licklider LJ, Gygi SP. Evaluation of Multidimensional Chromatography Coupled with Tandem Mass Spectrometry (LC/LC−MS/MS) for Large-Scale Protein Analysis:L The Yeast Proteome. J Proteome Res. 2003;2(1):43–50. doi:10.1021/pr025556v

132. Li W, Godzik A. Cd-hit: a fast program for clustering and comparing large sets of protein or nucleotide sequences. Bioinforma Oxf Engl. 2006;22(13):1658–1659. doi:10.1093/bioinformatics/btl158

133. Park SK, Venable JD, Xu T, Yates JR. A quantitative analysis software tool for mass spectrometry–based proteomics. Nat Methods. 2008;5(4):319–322. doi:10.1038/nmeth.1195

134. Huber W, von Heydebreck A, Sültmann H, Poustka A, Vingron M. Variance stabilization applied to microarray data calibration and to the quantification of differential expression. Bioinforma Oxf Engl. 2002;18 Suppl 1:S96–104. doi:10.1093/bioinformatics/18.suppl_1.s96

135. Gatto L, Lilley KS. MSnbase-an R/Bioconductor package for isobaric tagged mass spectrometry data visualization, processing and quantitation. Bioinforma Oxf Engl. 2012;28(2):288–289. doi:10.1093/bioinformatics/btr645

136. Ritchie ME, Phipson B, Wu D, et al. limma powers differential expression analyses for RNA-sequencing and microarray studies. Nucleic Acids Res. 2015;43(7):e47–e47. doi:10.1093/nar/gkv007

137. Gurdeep Singh R, Tanca A, Palomba A, et al. Unipept 4.0: Functional Analysis of Metaproteome Data. J Proteome Res. 2019;18(2):606–615. doi:10.1021/acs.jproteome.8b00716

138. Mesuere B, Debyser G, Aerts M, Devreese B, Vandamme P, Dawyndt P. The Unipept metaproteomics analysis pipeline. Proteomics. 2015;15(8):1437–1442. doi:10.1002/pmic.201400361

139. Sundararajan Z, Knoll R, Hombach P, Becker M, Schultze JL, Ulas T. Shiny-Seq: advanced guided transcriptome analysis. BMC Res Notes. 2019;12(1):432. doi:10.1186/s13104-019-4471-1

