## Supplemental Figures for "Peripheral Neuronal Activation Shapes the Microbiome and Alters Gut Physiology"

Figure S1

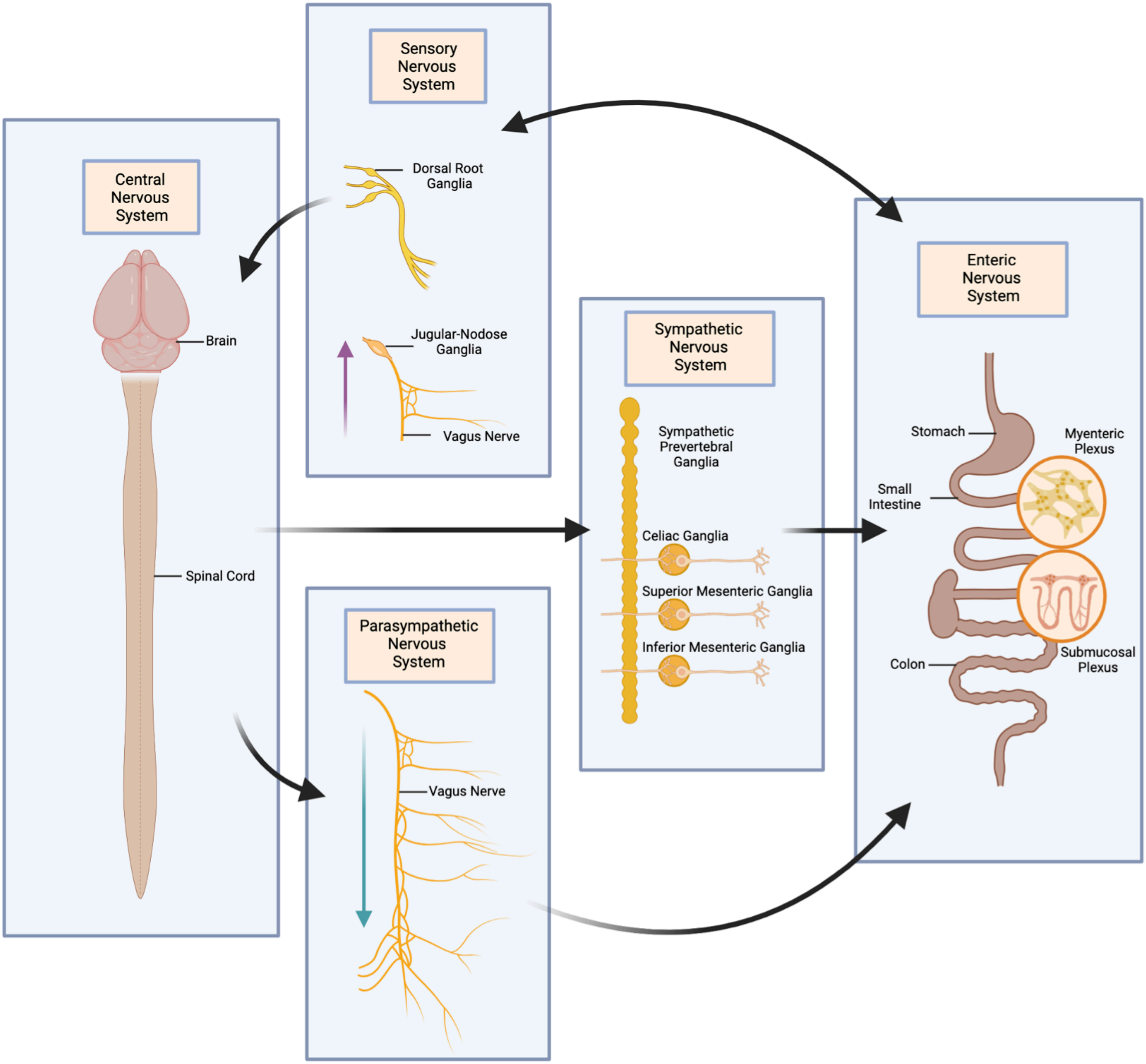

Sensory Nervous System
Dorsal Root Ganglia
Jugular-Nodose Ganglia
Vagus Nerve
Central Nervous System
Brain
Spinal Cord
Sympathetic Nervous System
Sympathetic Prevertebral Ganglia
Celiac Ganglia
Superior Mesenteric Ganglia
Inferior Mesenteric Ganglia
Enteric Nervous System
Stomach
Myenteric Plexus
Small Intestine
Submucosal Plexus
Colon
Parasympathetic Nervous System
Vagus Nerve

**Figure S1. Extrinsic and Intrinsic Innervation of the GI Tract (Related to Figure 1)**
Neurons of the sympathetic, parasympathetic, and sensory nervous systems extrinsically innervate the gut and have effects on GI function. In the sensory nervous system, sensory information from the gut is transmitted to the dorsal root ganglia (DRGs) and then to the spinal cord. Additionally, projections from the DRGs transmit sensory information to the gut. Other sensory signals arise from the periphery and are transmitted through the vagus nerve into the jugular-nodose ganglia and then into the brain. In the sympathetic nervous system, neurons from the spinal cord synapse onto neurons in the sympathetic prevertebral ganglia, celiac ganglia, superior mesenteric ganglia, and inferior mesenteric ganglia. These neurons then project to the gut. In the parasympathetic nervous system, the vagus nerve transmits signals from the brain to the gut. The intrinsic neurons of the gut, i.e., the enteric nervous system (ENS), reside in the myenteric and submucosal plexuses along the gut. Figure created with BioRender.com.

Figure S2

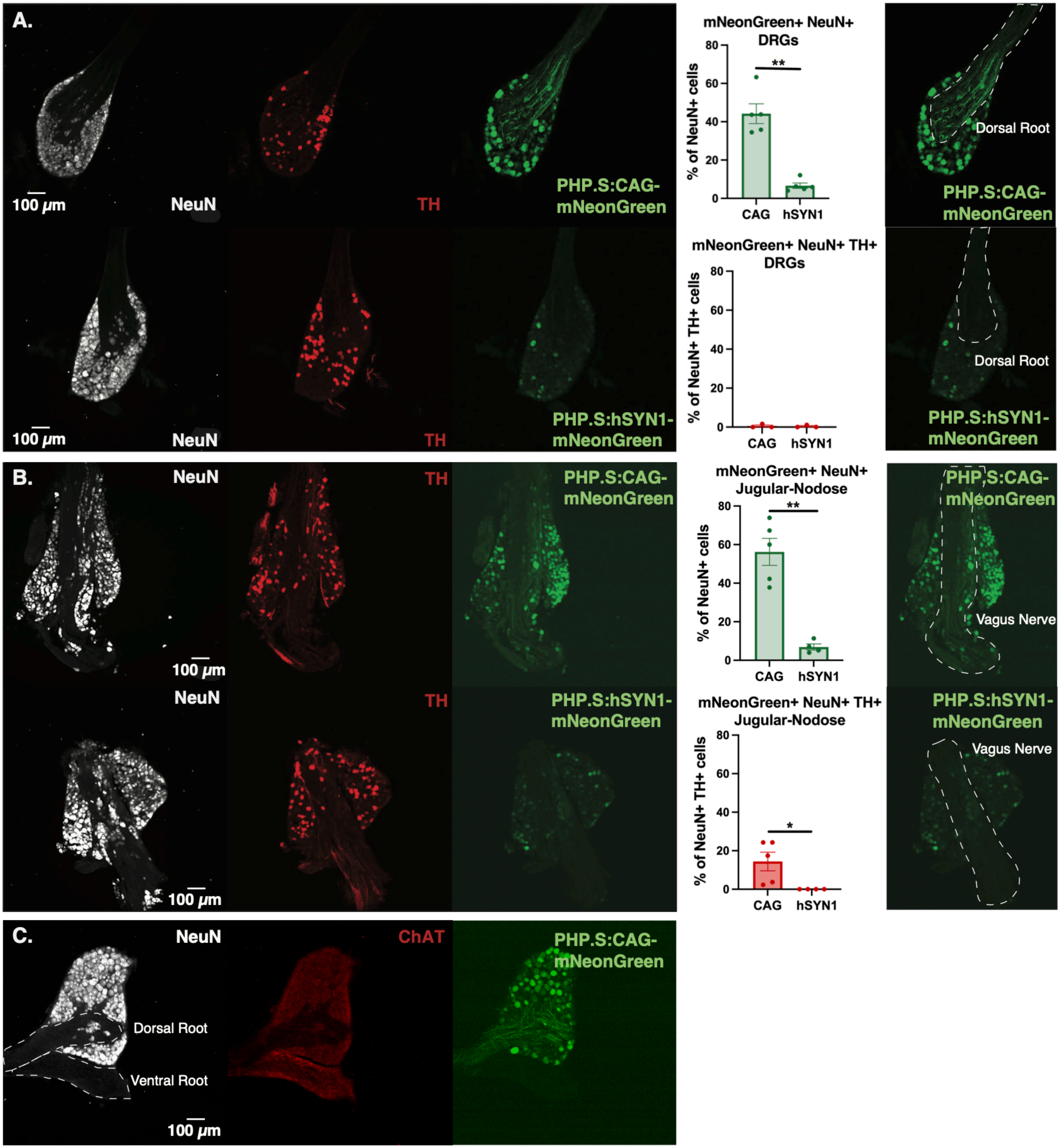
A.
100 μm
NeuN
TH
PHP.S:CAG-mNeonGreen
mNeonGreen+ NeuN+ DRGs
% of NeuN+ cells
80
60
40
20
0
**
CAG
hSYN1
Dorsal Root
PHP.S:hSYN1-mNeonGreen
mNeonGreen+ NeuN+ TH+ DRGs
% of NeuN+ TH+ cells
B.
mNeonGreen+ NeuN+ Jugular-Nodose
Vagus Nerve
mNeonGreen+ NeuN+ TH+ Jugular-Nodose
*
C.
ChAT
Dorsal Root
Ventral Root

**Figure S2. Viral Transduction of the Peripheral Nervous System (PNS) (Related to Figure 1)**

(A) Representative images showing labelling of the dorsal root ganglia (DRGs) by a single injection ($10^{12}$ vg) of either PHP.S-CAG-mNeonGreen or PHP.S-hSYN1-mNeonGreen. Cryosections were immunostained for the neuronal marker NeuN and tyrosine hydroxylase (TH). Plots show quantification of NeuN+ cells and NeuN+ TH+ cells virally labelled by AAV-PHP.S-CAG vs. AAV-PHP.S-hSyn1 in DRGs. (N=3-5 mice, each data point represents a 40 μM medial cross-section of the DRG, **$p<0.01$, determined by Welch's two-tailed t-test)

(B) Representative images showing labelling of the jugular-nodose ganglia by a single injection ($10^{12}$ vg) of either PHP.S-CAG-mNeonGreen or PHP.S-hSYN1-mNeonGreen. Cryosections were immunostained for the neuronal marker NeuN and TH. Plots show quantification of NeuN+ cells and NeuN+ TH+ cells virally labelled by AAV-PHP.S-CAG vs. AAV-PHP.S-hSyn1 in the jugular-nodose ganglia (N=4-5 mice, each data point represents a 40 μM medial cross-section of the jugular-nodose ganglia, **$p<0.01$, determined by Welch's two-tailed t-test).

(C) Representative images showing labelling of the dorsal root ganglia (DRGs), dorsal, and ventral root by a single injection ($10^{12}$ vg) of PHP.S-CAG-mNeonGreen. Cryosections were immunostained for the neuronal marker NeuN and ChAT. ChAT staining shows expected ChAT+ projections in the ventral root, but not the dorsal root or intrinsic neurons of the DRGs.

Figure S3

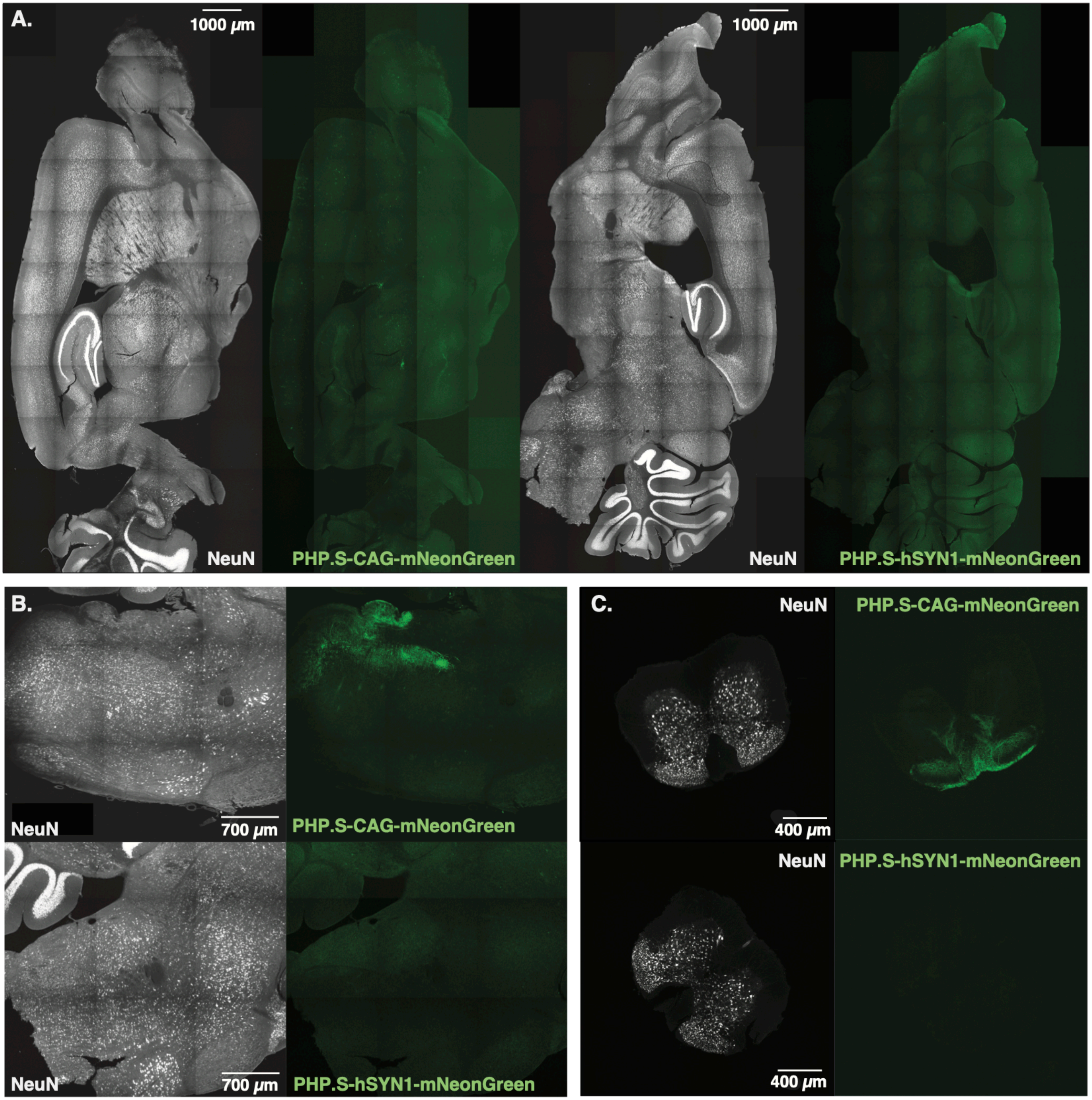
A.
1000 μm
1000 μm
NeuN
PHP.S-CAG-mNeonGreen
NeuN
PHP.S-hSYN1-mNeonGreen
B.
NeuN
700 μm
PHP.S-CAG-mNeonGreen
NeuN
700 μm
PHP.S-hSYN1-mNeonGreen
C.
NeuN
PHP.S-CAG-mNeonGreen
400 μm
NeuN
PHP.S-hSYN1-mNeonGreen
400 μm

#### Figure S3. Viral Transduction of the CNS by AAV-PHP.S (Related to Figure 1)

(A) Representative images showing labelling of the brain by a single injection ($10^{12}$ vg) of either AAV-PHP.S-CAG-mNeonGreen or AAV-PHP.S-hSYN1-mNeonGreen. Sections were immunostained for the neuronal marker NeuN.

(B, C) The PHP.S-hSYN1 injected mice did not show detectable mNeonGreen+ projections in the (B) brainstem or (C) spinal cord as were seen in the PHP.S-CAG injected mice.

Figure S4

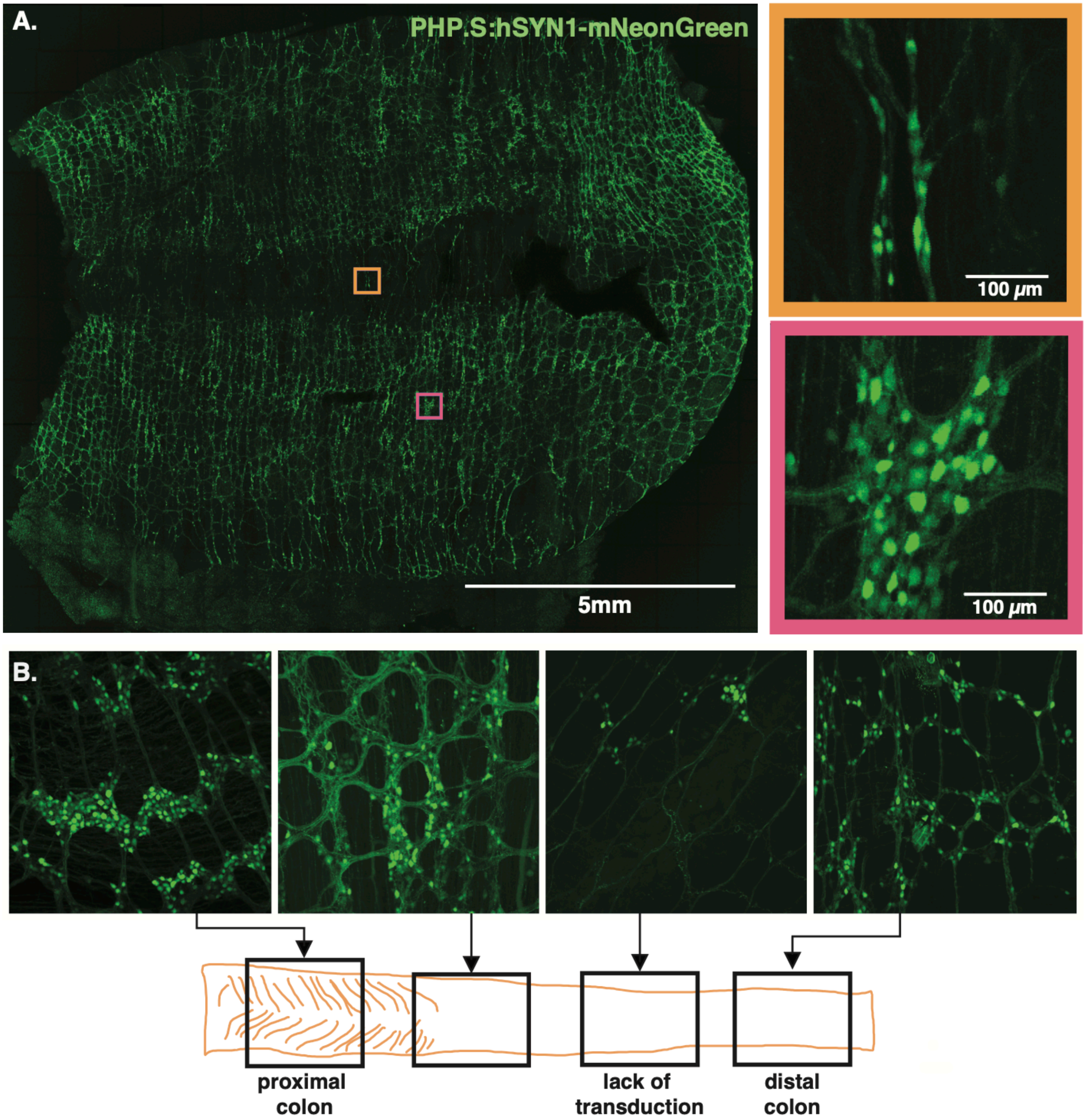

A.
PHP.S:hSYN1-mNeonGreen
5mm
100 μm
100 μm
B.
proximal colon
lack of transduction
distal colon

**Figure S4. Viral Transduction by AAV-PHP.S in the Colon (Related to Figure 1)**
(A) Tiled, whole mount microscopy showing widespread labelling in the proximal colon (>1 cm of tissue) from a single injection of AAV-PHP.S-hSYN1-mNeonGreen ($10^{12}$ vg). Orange inset highlights a region opposite the mesenteric attachment, and pink inset highlights a central region of the tissue.
(B) Representative images showing labelling patterns along the colon, including a 1.5 cm region that AAV-PHP.S does not transduce well.

### Figure S5

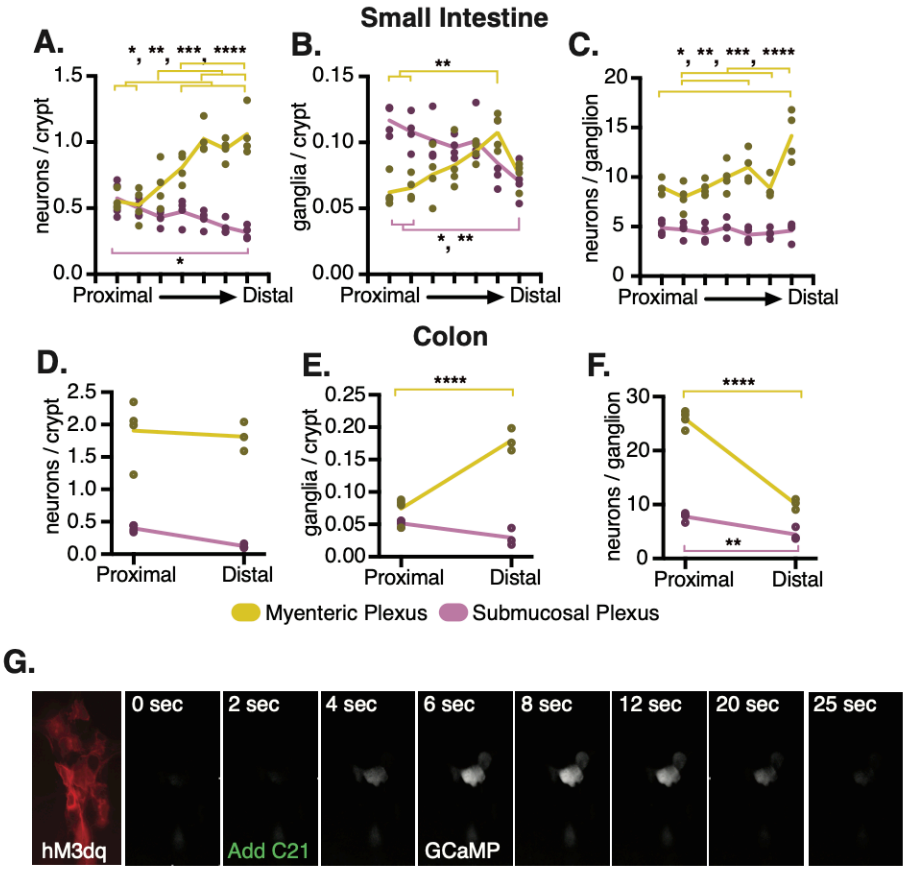

Small Intestine
A.
B.
C.
neurons / crypt
ganglia / crypt
neurons / ganglion
Proximal
Distal
Colon
D.
E.
F.
Myenteric Plexus
Submucosal Plexus
G.
hM3dq
0 sec
2 sec
Add C21
4 sec
6 sec
GCaMP
8 sec
12 sec
20 sec
25 sec

#### Figure S5. ENS Architecture and ChAT$^{+}$ Neuronal Stimulation in the ENS (Related to Figure 1)

(A-C) Quantification of neurons in 7 regions from the SI
(D-F) Quantification of neurons in 2 regions from the colon
(In A-F: N=3-4 mice per group, per intestinal region. Data points are averages from 2-3 images per mouse, per region; *: $p<0.05$, **: $p<0.01$, ***: $p<0.001$, ****: $p<0.0001$, determined by 2-way ANOVA with Sidak's multiple comparisons test)
(G) Fluorescence time course of GCaMP6f following C21 administration in an *ex vivo* preparation of ChAT$^{+}$ SI injected with PHP.S-hSYN1-DIO-hM3Dq-mRuby2 and PHP.S-CAG-GCaMP6f.

### Figure S6

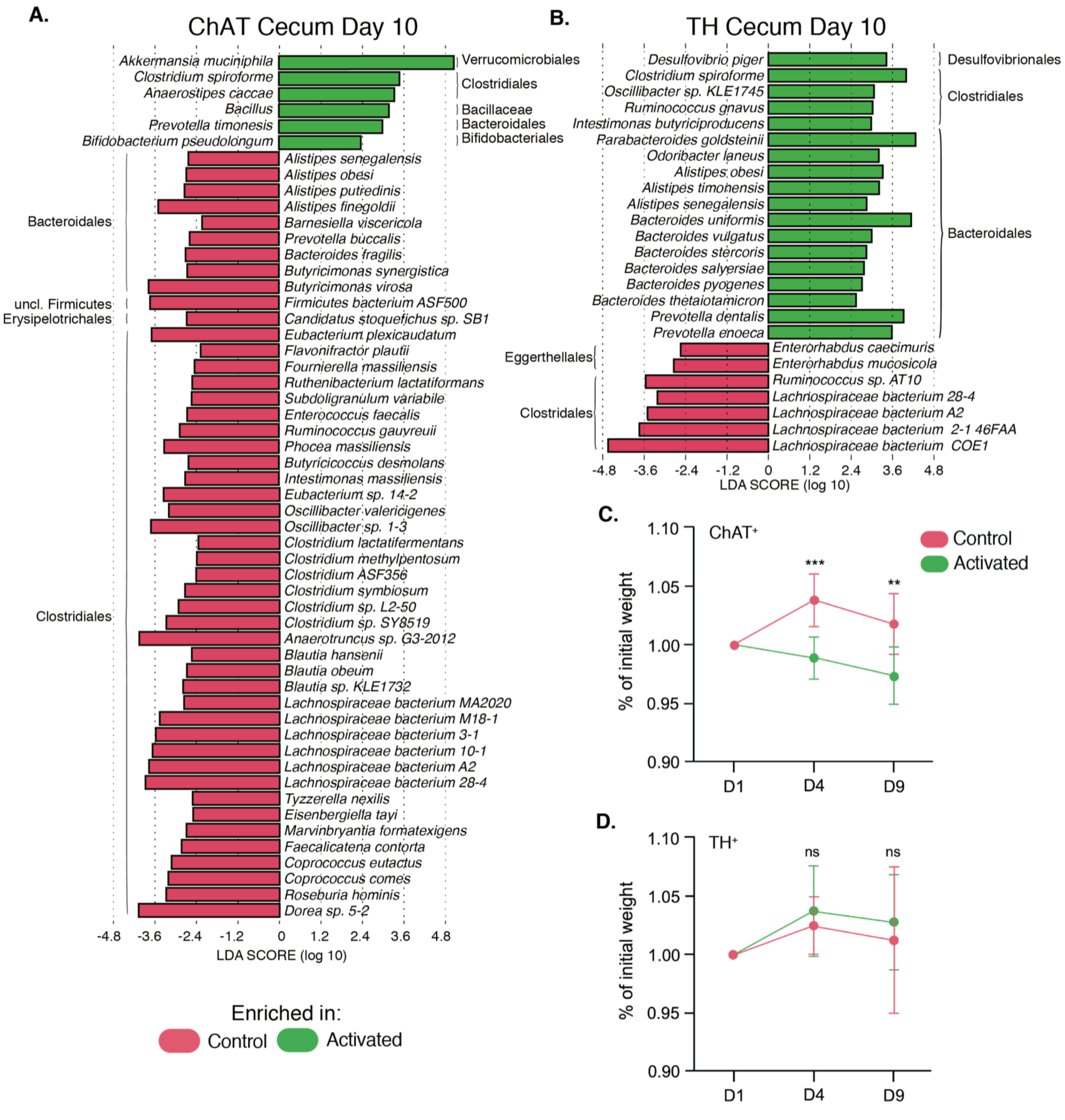

A.
ChAT Cecum Day 10
Akkermansia muciniphila
Clostridium spiroforme
Anaerostipes caccae
Bacillus
Prevotella timonensis
Bifidobacterium pseudolongum
Verrucomicrobiales
Clostridiales
Bacillaceae
Bacteroidales
Bifidobacteriales
Alistipes senegalensis
Alistipes obesi
Alistipes putredinis
Alistipes finegoldii
Barnesiella viscericola
Prevotella buccalis
Bacteroides fragilis
Butyricimonas synergistica
Butyricimonas virosa
Firmicutes bacterium ASF500
Candidatus stoquefichus sp. SB1
Eubacterium plexicaudatum
Flavonifractor plautii
Fournierella massiliensis
Ruthenibacterium lactatiformans
Subdoligranulum variabile
Enterococcus faecalis
Ruminococcus gauvreuii
Phocea massiliensis
Butyricicoccus desmolans
Intestimonas massiliensis
Eubacterium sp. 14-2
Oscillibacter valericigenes
Oscillibacter sp. 1-3
Clostridium lactatifermentans
Clostridium methylpentosum
Clostridium ASF356
Clostridium symbiosum
Clostridium sp. L2-50
Clostridium sp. SY8519
Anaerotruncus sp. G3-2012
Blautia hansenii
Blautia obeum
Blautia sp. KLE1732
Lachnospiraceae bacterium MA2020
Lachnospiraceae bacterium M18-1
Lachnospiraceae bacterium 3-1
Lachnospiraceae bacterium 10-1
Lachnospiraceae bacterium A2
Lachnospiraceae bacterium 28-4
Tyzzerella nexilis
Eisenbergiella tayi
Marvinbryantia formatexigens
Faecalicatena contorta
Coprococcus eutactus
Coprococcus comes
Roseburia hominis
Dorea sp. 5-2
Bacteroidales
uncl. Firmicutes
Erysipelotrichales
Clostridiales
-4.8 -3.6 -2.4 -1.2 0 1.2 2.4 3.6 4.8
LDA SCORE (log 10)
Enriched in:
Control
Activated
B.
TH Cecum Day 10
Desulfovibrio piger
Clostridium spiroforme
Oscillibacter sp. KLE1745
Ruminococcus gnavus
Intestimonas butyriciproducens
Parabacteroides goldsteinii
Odoribacter laneus
Alistipes obesi
Alistipes timonensis
Alistipes senegalensis
Bacteroides uniformis
Bacteroides vulgatus
Bacteroides stercoris
Bacteroides salyersiae
Bacteroides pyogenes
Bacteroides thetaiotamicron
Prevotella dentalis
Prevotella enoeca
Desulfovibrionales
Clostridiales
Bacteroidales
Eggerthellales
Clostridales
Enterorhabdus caecimuris
Enterorhabdus mucosicola
Ruminococcus sp. AT10
Lachnospiraceae bacterium 28-4
Lachnospiraceae bacterium A2
Lachnospiraceae bacterium 2-1 46FAA
Lachnospiraceae bacterium COE1
-4.8 -3.6 -2.4 -1.2 0 1.2 2.4 3.6 4.8
LDA SCORE (log 10)
C.
ChAT+
Control
Activated
***
**
% of initial weight
1.10
1.05
1.00
0.95
0.90
D1
D4
D9
D.
TH+
ns
ns
% of initial weight
1.10
1.05
1.00
0.95
0.90
D1
D4
D9

**Figure S6. Annotated changes in gut bacteria (Related to Figure 2 and Figure 6)**
(A,B) Species-levels differences in cecal bacteria after 10 days of C21 injection in (A) $ChAT^{+}$ and (B) $TH^{+}$ mice
(C,D) Analysis of activation-mediated changes in body weight of (C) $ChAT^{+}$ and (D) $TH^{+}$ mice over 9 days of C21 administration
(N = 10-11 mice per group; **: p<0.01, ***: p<0.001, determined by 2-way ANOVA with Sidak's method for multiple comparisons)

### Figure S7

A.

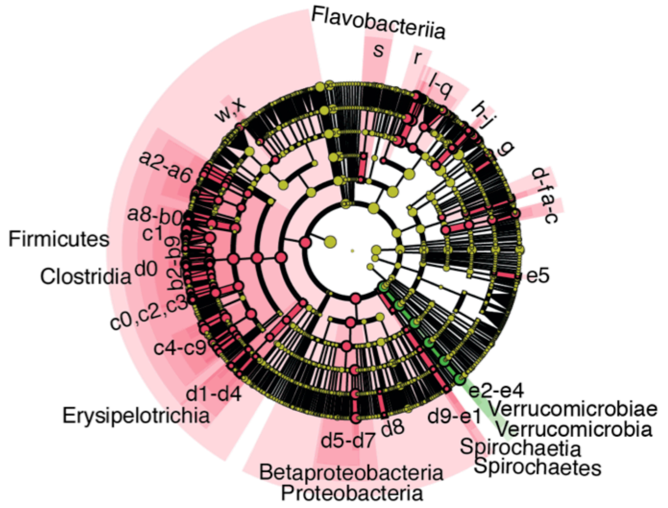


B.

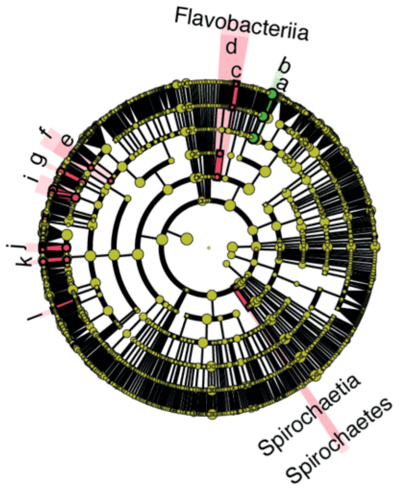


| ChAT-Cre Day 10 Feces | | | | | | | |
|---|---|---|---|---|---|---|---|
| **ID** | **Microbial Taxon** | **ID** | **Microbial Taxon** | **ID** | **Microbial Taxon** | **ID** | **Microbial Taxon** |
| a | Bifidobacterium | p | Parabacteroides | b4 | Dorea | d0 | Clostridiales |
| b | Bifidobacteriaceae | q | Tannerellaceae | b5 | Eisenbergiella | d1 | Candidatus Stoquefichus |
| c | Bifidobacteriales | r | Marinilbiliales | b6 | Faecalicatena | d2 | Faecalitalea |
| d | Cellulomonas | s | Flavobacteriales | b7 | Lachnoclostridium | d3 | Erysipelotrichaceae Lorem ipsum |
| e | Cellulomondaceae | w | Enterococcus | b8 | Marvinbryantia | d4 | Erysipelotrichales |
| f | Micrococcales | x | Enterococcaceae | b9 | Sellimonas | d5 | Parasutterella |
| g | Adlercreutzia | a2 | Intestinimonas | c0 | Tyzzerella | d6 | Sutterellaceae |
| h | Barnsiella | a3 | Pseudoflavonifractor | c1 | Lachnospiraceae | d7 | Burkholderiales |
| i | Coprobacter | a5 | Clostridium | c2 | Oscillibacter | d8 | Lawsonia |
| j | Barnesiellaceae | a6 | Clostridiaceae | c3 | Oscillospiraceae | d9 | Treponema |
| k | Prevotella | a8 | Anaerofustis | c5 | Anaerotruncus | e0 | Spirochaetaceae |
| l | Prevotellaceae | a9 | Eubacterium | c6 | Ruminiclostridium | e1 | Spirochaetales |
| m | Alistipes | b0 | Eubacteriaceae | c7 | Ruminococcus | e2 | Akkermansia |
| n | Rikenella | b2 | Blautia | c8 | Ruthenibacterium | e3 | Akkermansiaceae |
| o | Rikenellaceae | b3 | Coprococcus | c9 | Ruminococcaceae | e4 | Verrucomicrobiales |
| | | | | | | e5 | Varicellovirus |

| TH-Cre Day 6 Feces | |
|---|---|
| **ID** | **Microbial Taxon** |
| a | Parabacteroides |
| b | Tannerellaceae |
| c | Fluviicola |
| d | Flavobacteriales |
| e | Streptococcus |
| f | Streptococcaceae |
| g | Pseudoflavonifractor |
| i | Clostridium |
| j | Blautia |
| k | Eisenbergiella |
| l | Clostridiodes |

Enriched in:
Control
Activated

C.

| ChAT-Cre Cecum | | |
|---|---|---|
| **ID** | **Gene Family** | **Annotation** |
| 1 | K02003 | putative ABC transport system ATP-binding protein |
| 2 | K02518 | translation initiation factor IF-1 |
| 3 | K02948 | small subunit ribosomal protein S11 |
| 4 | K02950 | small subunit ribosomal protein S12 |
| 5 | K02965 | small subunit ribosomal protein S19 |
| 6 | K03205 | type IV secretion system protein VirD4 |
| 7 | K03294 | basic amino acid/polyamine antiporter, APA family |
| 8 | K03496 | chromosome partitioning protein |
| **ID** | **Gene Pathway** | **Annotation** |
| 1 | HISDEG-PWY | L-histidine degradation I |
| 2 | ILEUSYN-PWY | L-isoleucine biosynthesis I (from threonine) |
| 3 | PWY-6122 | 5-aminoimidazole ribonucleotide biosynthesis II |
| 4 | PWY-6595 | superpathway of guanosine nucleotides degradation (plants) |
| 5 | PWY-6606 | guanosine nucleotides degradation II |
| 6 | PWY-7111 | pyruvate fermentation to isobutanol (engineered) |
| 7 | PWY-7219 | adenosine ribonucleotides de novo biosynthesis |
| 8 | VALSYN-PWY | L-valine biosynthesis |

D.

| ChAT-Cre Day 10 Feces | | |
|---|---|---|
| **ID** | **Gene Family** | **Annotation** |
| 1 | K02003 | putative ABC transport system ATP-binding protein |
| 2 | K02518 | translation initiation factor IF-1 |
| 3 | K02913 | large subunit ribosomal protein L33 |
| 4 | K02946 | small subunit ribosomal protein S10 |
| 5 | K02948 | small subunit ribosomal protein S11 |
| 6 | K02950 | small subunit ribosomal protein S12 |
| 7 | K02965 | small subunit ribosomal protein S19 |
| 8 | K03205 | type IV secretion system protein VirD4 |
| **ID** | **Gene Pathway** | **Annotation** |
| 1 | ILEUSYN-PWY | L-isoleucine biosynthesis I (from threonine) |
| 2 | PWY-6151 | S-adenosyl-L-methionine cycle I |
| 3 | PWY-6700 | queuosine biosynthesis I (de novo) |
| 4 | PWY-6859 | all-trans-farnesol biosynthesis |
| 5 | PWY-7111 | pyruvate fermentation to isobutanol |
| 6 | PWY-7219 | adenosine ribonucleotides de novo biosynthesis |
| 7 | PWY0-1296 | purine ribonucleosides degradation |
| 8 | VALSYN-PWY | L-valine biosynthesis |

#### Figure S7. Gut-Associated Activation-Mediated Changes in the Fecal Microbiome (Related to Figure 2)

(A) LEfSe cladograms and annotations for feces on Day 10 of C21 administration in $ChAT^{+}$ mice
(B) LEfSe Cladograms and annotations for feces on Day 6 of C21 administration in $TH^{+}$ mice
(C-D) Microbial gene family and pathway annotations for the feature arrows in Figures 2P-2S. Colors in the ID column refer to the colors of the arrows in the main figure.
